# Fungal antibiotics control bacterial community diversity in the cheese rind microbiome

**DOI:** 10.1101/2022.11.26.518062

**Authors:** Joanna Tannous, Casey M. Cosetta, Milton T. Drott, Tomás A. Rush, Paul E. Abraham, Richard J. Giannone, Nancy P. Keller, Benjamin E. Wolfe

**Affiliations:** Department of Medical Microbiology and Immunology, University of Wisconsin – Madison, Madison, WI, USA; Biosciences Division, Oak Ridge National Laboratory, Oak Ridge, TN 37831, USA; Department of Biology, Tufts University, Medford, MA, USA; Department of Plant Pathology, University of Wisconsin-Madison, Madison, WI, USA; USDA-ARS Cereal Disease Laboratory, St. Paul, MN 55108, USA

## Abstract

Potent antimicrobial metabolites are produced by filamentous fungi in pure lab cultures, but their ecological functions in nature are often unknown. Using an antibiotic-producing *Penicillium* isolate and the cheese rind microbial community, we demonstrate that a fungal specialized metabolite can regulate the diversity of bacterial communities. Inactivation of the global regulator, LaeA, resulted in the loss of antibacterial activity in the *Penicillium* isolate. Cheese rind bacterial communities assembled with the *laeA* deletion strain had significantly higher bacterial abundances than the wild-type strain. RNA-sequencing and metabolite profiling demonstrated a striking reduction in the expression and production of the natural product pseurotin in the *laeA* deletion strain. Inactivation of a core gene in the pseurotin biosynthetic cluster restored bacterial community composition, demonstrating the role of pseurotins in mediating bacterial community assembly. Our discovery demonstrates how antibiotic production can drive the assembly of microbiomes and provides an ecological context for widespread fungal specialized metabolites.

## INTRODUCTION

Fungal specialized (a.k.a secondary) metabolites (SMs) gained attention in the 1900s following the discovery of the world’s first antibiotic, penicillin, produced by a *Penicillium* isolate^1^. Since then, fungal SMs have come to play pivotal roles in medicine, agriculture, and biotechnology. Many aspects of fungal SMs have been well-characterized in axenic cultures, including genetic regulation^2–5^, biochemistry^6^, and impacts on the physiology and development of their producers^7–9^. In co-cultures, fungal SMs have been suggested to play roles as signaling molecules that mediate the communication of the fungus with its surroundings^10–13^, virulence factors to support pathogenic lifestyles^14, 15^, microbial inhibitors that shape the competition with other microorganisms for finite resources^16–18^, or armaments against fungivores^19, 20^.

While these studies are critical first steps in understanding the potential functions of fungal SMs, the ecological roles of these compounds in multispecies communities are largely unknown. The activities of fungal SMs identified under laboratory conditions may not translate to natural communities if concentrations used *in vitro* are not reflective of those found in nature or if other community members are able to inactivate or alter these compounds. Purified and concentrated fungal SMs can inhibit the growth of some bacteria and fungi in large quantities in simplified lab environments^21, 22^, but how naturally secreted fungal SMs operate in microbial communities is unknown. Fungal SMs could structure multispecies communities by mediating ecological interactions that favor certain species over others, resulting in a shift in community composition. We are unaware of studies that have demonstrated this scenario.

One approach for identifying fungal SMs that could mediate bacterial communities is by altering the activity of global regulators of fungal metabolites. In the fungal phylum Ascomycota, the production of many SMs is under the control of the global regulatory protein of the trimeric velvet complex, LaeA^23, 24^. A growing body of research has emphasized the role of LaeA in regulating SM production in monocultures of *Aspergillus*, *Fusarium*, *Penicillium*, and other fungal genera^25–28^. Several studies have explored how LaeA can mediate fungal strain competition and host-microbe interactions^8, 29^, but the ecological role of LaeA in the assembly of polymicrobial communities has not been characterized.

Cheese rinds are microbial ecosystems where LaeA-regulated SMs could have significant ecological impacts. These ecosystems are composed of bacteria, yeasts, and filamentous fungi and form on the surface of many styles of cheese, including bloomy, washed, and natural rind cheeses^30^. *Penicillium* sp. are frequently encountered in cheese rinds where they can be inoculated as industrial starter cultures (e.g. *P. camemberti* in Camembert or Brie) or can colonize cheese from natural populations of these fungi (e.g. *P. biforme, P. solitum,* and *P. nalgiovense* in tomme style cheeses and clothbound Cheddars^30, 31^). Species within this genus are prolific producers of SMs, including polyketides (e.g. patulin), non-ribosomal peptides (e.g. roquefortine) and terpenes (e.g. expansolide)^32^. While many *Penicillium* metabolites are valued as pharmaceuticals, such as the antibiotic penicillin^1^ and the cholesterol-lowering drug lovastatin^33^, others are considered mycotoxins, including the carcinogenic ochratoxin A^34^, cyclopiazonic acid^35^, and patulin^36^. *Penicillium* species isolated from cheese rinds produce an extensive range of SMs, including mycotoxins^37, 38^, and have been shown to have a strong impact on the growth of neighboring bacteria^39–41^, suggesting a potential of fungi to control bacterial community diversity through antibiotic production.

In the current study, we determined the ecological significance of fungal SMs in the cheese rind model system by inactivating LaeA in a cheese rind *Penicillium* strain (*Penicillium* sp. strain MB). This fungus is closely related to *Penicillium polonicum* ^42^ and was disrupting rind development when it was originally isolated from a natural-rind cheese. Given its negative impacts on the cheese rind, we predicted that this strain produced secondary metabolites that could alter rind community assembly. When we deleted *laeA* in this strain, an *in vitro* cheese rind bacterial community increased in total abundance and shifted in composition to resemble bacterial communities grown without the fungus. Both transcriptomic analysis and metabolite profiling pointed to pseurotins as putative LaeA-regulated antibacterial compounds. Inactivation of pseurotin production in the WT strain through the disruption of the gene encoding the hybrid PKS-NRPS enzyme required for pseurotin synthesis eliminated much of the antibacterial activity and caused a shift in bacterial community composition that was similar to the Δ*laeA* strain. This work demonstrates the ecological relevance of fungal SMs, their roles in shaping the assembly of multispecies bacterial communities, and their possible influence on the development of human food commodities.

## RESULTS

### Confirmation of targeted gene edits in *Penicillium sp.* strain MB

Confirmation of *laeA* deletion in *Penicillium* strain MB_WT— Twenty-five hygromycin-resistant transformants were isolated after a rapid selection procedure on SMM supplemented with hygromycin. About 40% (10/25) of the monoconidial lines generated from primary transformants of strain MB were PCR-positive for the absence of the *laeA* ORF (data not shown). Southern blotting using probes corresponding to either 5’or 3’ flanks of *laeA* (used in the construction of the deletion cassette) revealed single-site integration of the deletion cassette in a single transformant of the strain MB (**Supplementary Fig. 1b**). A HygR control strain, which has the hygromycin cassette inserted into the genome but not at the target locus, was also included in the study as a control for absence of selectable marker gene effects.

Confirmation of *laeA* complementation in *Penicillium* strain MB_Δ*laeA*— Ten phleomycin-resistant transformants were isolated and subjected to Southern blot to confirm the single insertion of the *laeA* ORF. One strain out of ten showed a single band of 2.8 Kb that matches the band obtained with the positive control (plasmid PJT3 generated after subcloning the *laeA* fragment into plasmid pBC-phleo). As expected, the MB_Δ*laeA* mutant strain used as a negative control did not show any band (**Supplementary Fig. 1d**).

Confirmation of *psoA* deletion in *Penicillium* strain MB_WT using Cas9/sgRNA RNP complex— Four hygromycin resistant transformants were isolated after a rapid selection procedure on SMM supplemented with hygromycin. Two out of four of the monoconidial lines generated were PCR-positive for the absence of the psoA ORF (data not shown). Later, the off-target events in the two CRISPR-Cas9-modified strains were assessed by whole-genome sequencing analysis using Illumina sequencing. Overall, only two high-confidence SNPs were detected in both CRISPR knockout strains that could have resulted from off-target effects: one SNP (G→A with predicted amino acid substitution of D→N) in a gene annotated as “chromatin structure-remodeling complex subunit rsc1” and another SNP (DNA G→A with predicted amino acid substitution of P→L) in a predicted protein with an unknown function.

### The deletion of *laeA* impairs several physiological traits in *Penicillium* sp. strain MB

The deletion and complementation of *laeA* in *Penicillium* strain MB resulted in four strains: MB_WT, MB_Δ*laeA*, MB_Hyg^R^ (to test for effects of inserting the hygromycin selectable marker into the genome) and MB_*laeA*c (complementation of the Δ*laeA* strain). The MB_Δ*laeA* strain had altered pigmentation and reduced spore production relative to MB_WT, and complementation of the Δ*laeA* strain (MB_*laeA*c) restored the WT phenotype (**Fig. 1a-b**). Differences in pigmentation were most striking on cheese curd agar (CCA), a medium that mimics cheese surface environments in cheese aging facilities^39, 43^. No differences in pigment or sporulation phenotypes were observed between the MB_WT, the MB_Hyg^R^, and the MB_*laeA*c strains.

**Figure 1:**
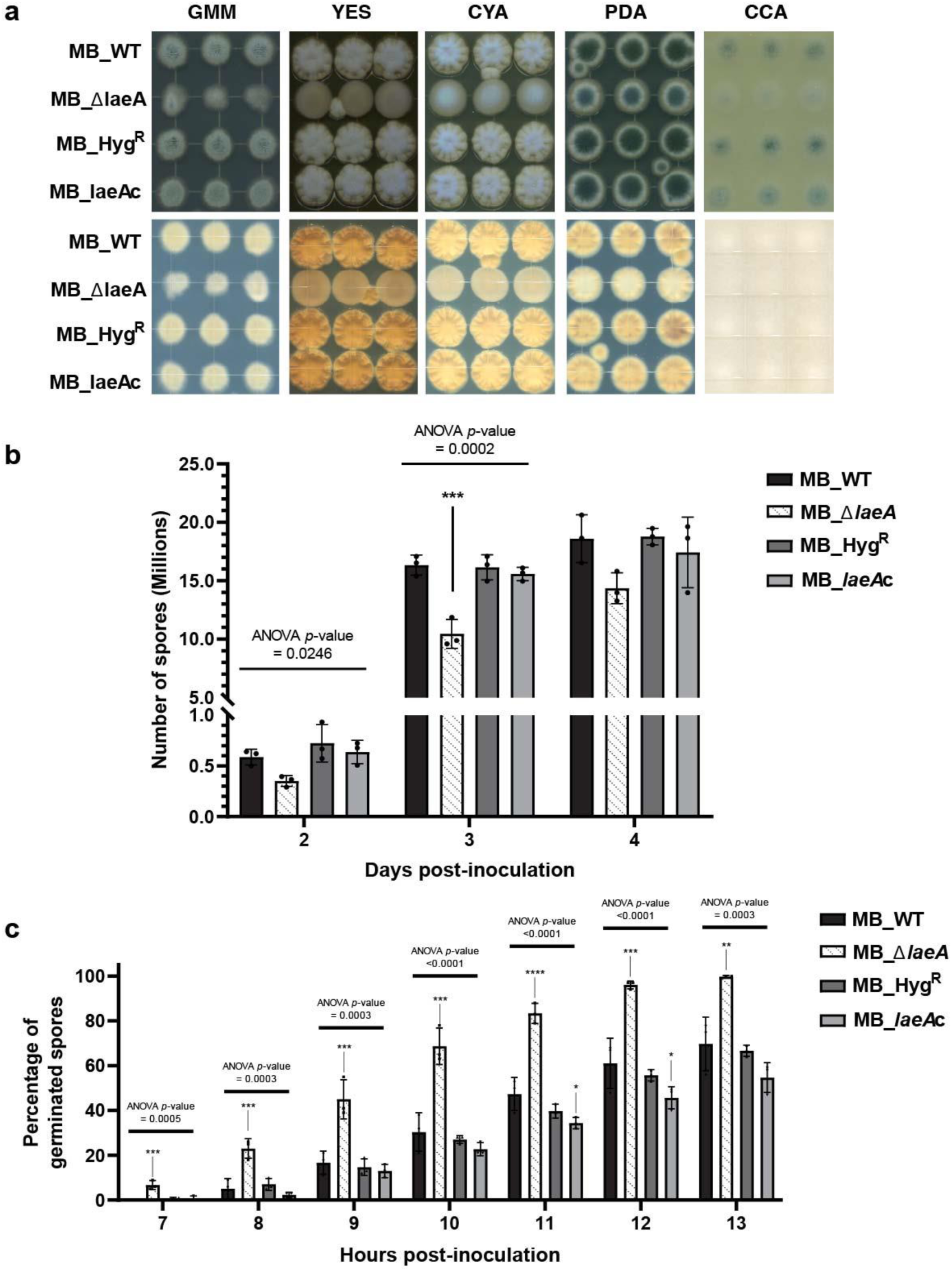
Deletion of *laeA* affects both morphological and physiological characteristics of *Penicillium* sp. MB strains. (a) Colony aspect of *Penicillium* sp. MB strains grown on different media for 5 days at 25 °C. GMM = glucose minimal medium, YES = yeast extract medium, CYA = Czapek Yeast Autolysate Agar, PDA = potato dextrose agar, CCA = cheese curd agar. (b) Spore count for each strain over four days growth on GMM agar medium (c) Percentage of germinated spores over 13 hours growth in GMM broth. Counting germinated spores started 3 hours post-inoculation. There were no germinated spores between hours 3 to 6. In the box plots, the bar represents the standard errors of the means and each dot represents a biological replication (n = 3). One-way analysis of variance (ANOVA) was performed for each day for the sporulation data and each hour for the germination data. Dunnett’s multiple comparison test was used and compared to the MB_WT strain. (****) indicates a *p*-value < 0.0001, (***) indicates a *p*-value < 0.001, (**) indicates a *p*-value < 0.01, (*) indicates a *p*-value < 0.05, and no asterisk indicates not significant. For exact *p*-values for each treatment, see **Dataset 1.**

The deletion of *laeA* resulted in an accelerated germination of spores, reaching 100% germination 13 hours post-inoculation, while none of the control strains were able to achieve complete spore germination (**Fig. 1c**). Many fungi produce self-inhibitors that reduce germination rates, especially at high inoculum levels, and it is possible that *laeA* loss results in the diminishment of these self-inhibitors ^44^. The deletion of *laeA* also resulted in reduced growth (as measured by colony diameter) regardless of the media **(Supplementary Fig. 2**). Collectively, these growth and development data demonstrate that LaeA regulates fungal traits that could have consequences for neighboring microbes.

### *Penicillium* traits regulated by LaeA mediate cheese rind bacterial community assembly and microbial interactions

To test how LaeA regulation of physiological or metabolic traits alters the development of cheese rind microbiomes, we grew all four *Penicillium* MB strains (MB_WT, MB_Δ*laeA*, MB_Hyg^R^ and MB_*laeA*c) with a four-member bacterial community that represents the dominant taxa found in typical natural rind cheeses. Here we compared total bacterial community abundance (as total CFUs) and bacterial community composition (relative abundance of each community member) at 3-, 10- and 21-days post-inoculation across five different treatments: **1)** bacteria alone, **2)** bacteria + *Penicillium* WT strain, **3)** bacteria + *Penicillium* Δ*laeA* strain (MB_Δ*laeA)*, **4)** bacteria + *Penicillium* hygromycin resistance control strain (MB_Hyg^R^), and **5)** bacteria + *Penicillium laeA* complement strain (MB_*laeA*c).

After 3 days of incubation, community composition was similar across all treatments (**Fig. 2a, b**). However, the presence of MB_WT, MB_Hyg^R^ and MB_*laeA*c strains caused a significant restructuring of the bacterial community through their strong inhibitory effects at day 10 post-inoculation (**Fig. 2c**). The bacteria alone treatments were dominated by Actinobacteria (*Brevibacterium* 30% and *Brachybacterium* 38%) whereas the MB_WT, MB_Hyg^R^ and MB_*laeA*c treatments were *Staphylococcus*-dominated (over 82% of the population) (**Fig. 2c**). In the presence of MB_WT, the absolute growth for all bacteria was significantly reduced, with *Brevibacterium* and *Brachybacterium* having nearly a 4-fold decrease in absolute growth, and *Psychrobacter* populations not reaching detectable levels (**Fig. 2d**). Comparable results were observed with the *Penicillium* MB_Hyg^R^ and MB_*laeA*c strains. Strikingly, the abundance and structure of the bacterial community in the presence of MB_Δ*laeA* was similar to the community grown in the absence of the fungus (a range of 57-68% Actinobacteria) (**Fig. 2c**). The shifts in community composition and dominance of Actinobacteria in the bacteria-alone and MB_Δ*laeA* communities persisted through day 21 (**Fig. 2e, f**).

**Figure 2:**
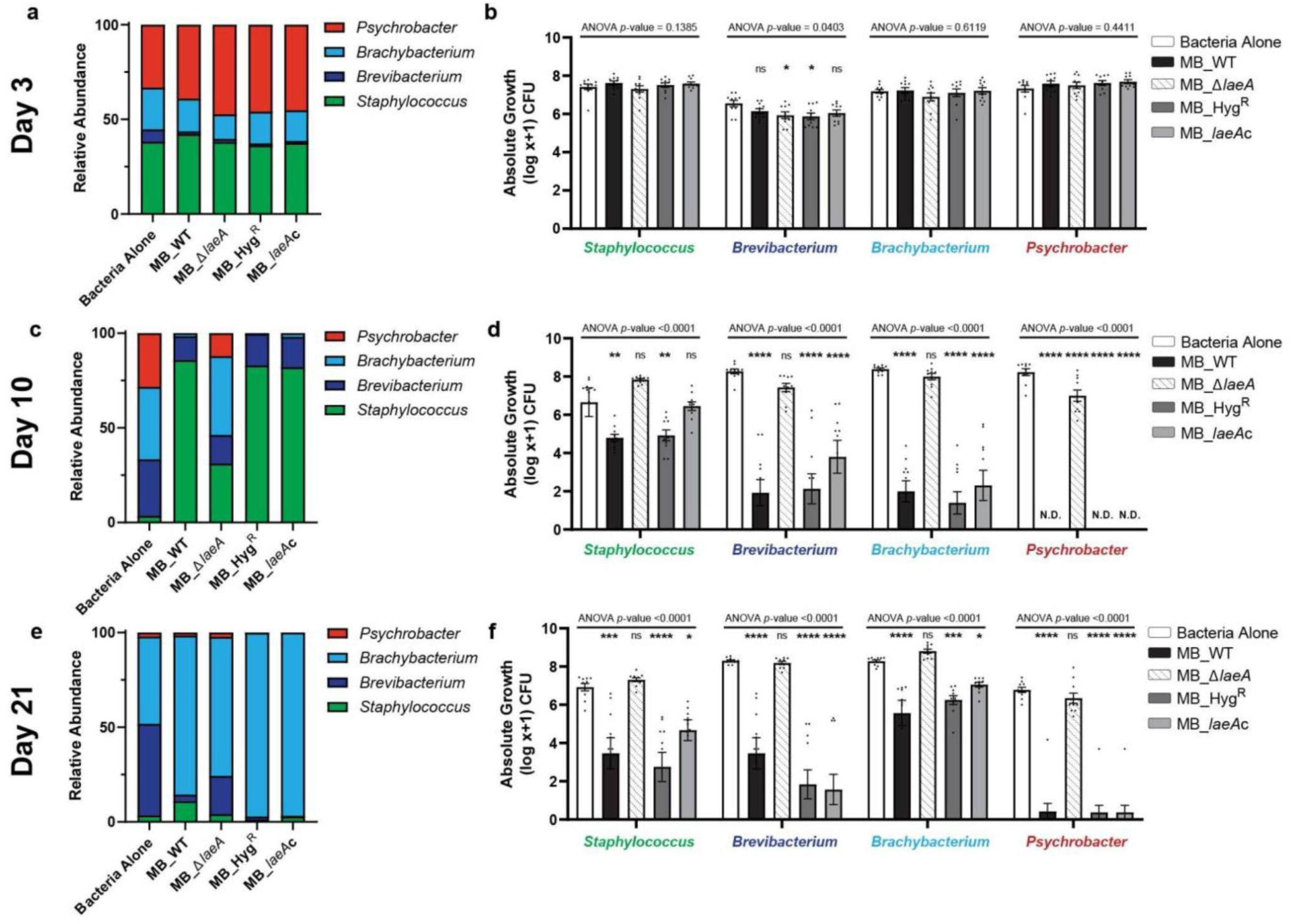
Inactivation of LaeA increases bacterial diversity and abundance in cheese rind communities. Histograms showing the shifts in a typical rind community (*Staphylococcus, Brevivacterium, Brachybacterium* and *Psychrobacter*) in the presence of four strains of *Penicillium* sp. strain MB after **(a) 3, (c) 10 and (e)** 21 days of incubation. Data are mean relative abundance from two independent experiments with five biological replications each. The compositions of the Bacteria Alone and MB_Δ*laeA* communities were not significantly different from one another, but were different from MB_WT, MB_Hyg^R^, MB_*laeAc* at 10 and 21 days (Day 3 PERMANOVA F = 0.9958 p = 0.425; Day 10 PERMANOVA F = 23.12, p < 0.0001; Day 21 PERMANOVA F = 11.14, p < 0.0001). Histograms showing the inhibition of community members grown in the presence of *Penicillium* sp. strain WT and mutants (MB_Δ*laeA*, MB_Hyg^R^, and MB_*laeAc*) after **(b)** 3 **(d)**, 10 and **(f)** 21 days of incubation. Each bar represents the mean with standard errors and each dot represents a biological replicate. One-way analysis of variance (ANOVA) was performed for each bacterium. Dunnett’s multiple comparison test was used and compared to the WT strain. (****) indicates a *p*-value < 0.0001, (***) indicates a *p*-value < 0.001, (**) indicates a *p*-value < 0.01, (*) indicates a *p*-value < 0.05, and no asterisk indicates not significant (ns). For exact *p*-values for each treatment, see **Dataset 1.**

**Supplementary Fig. 6:**
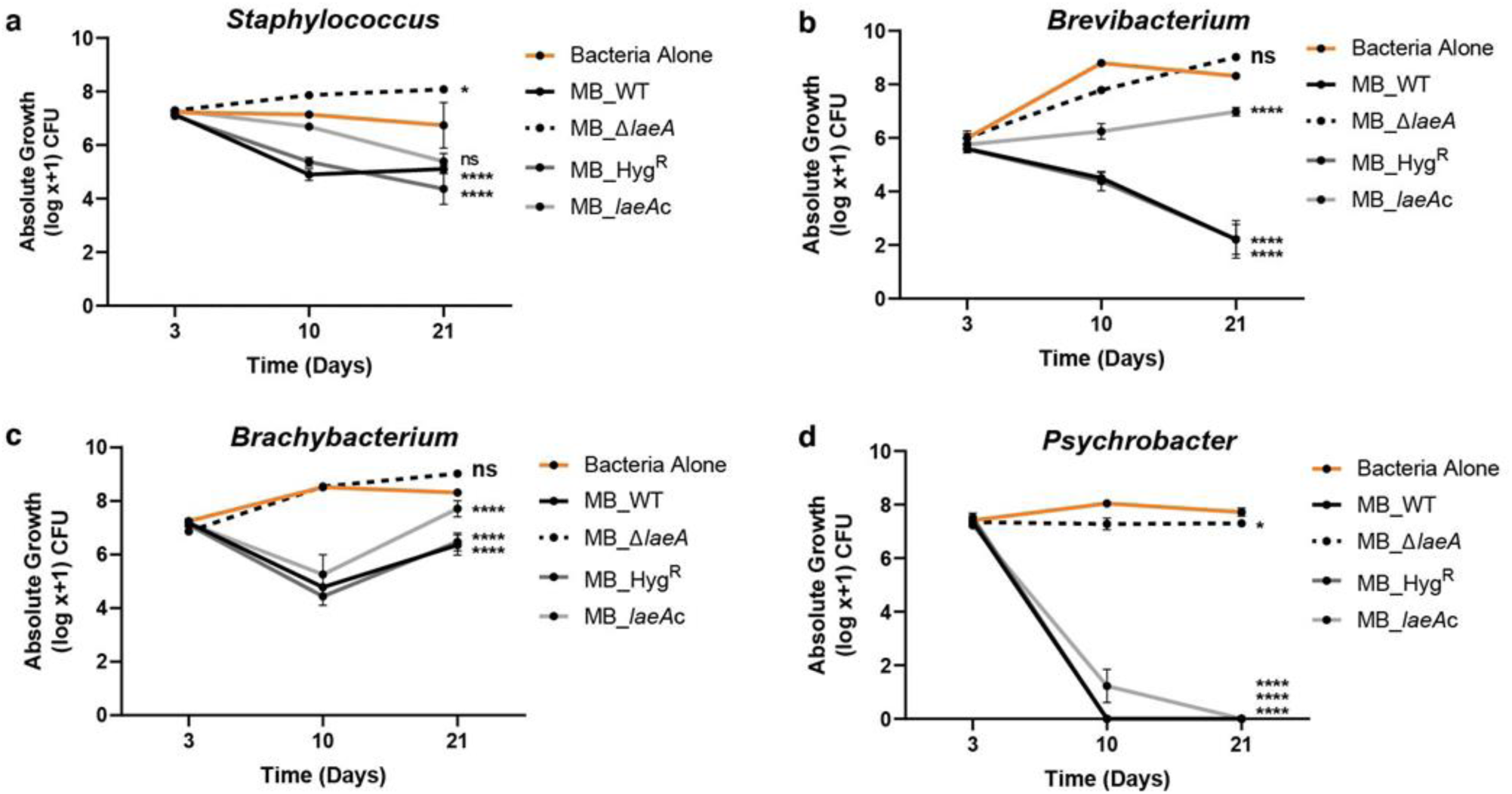
Pairwise interaction assays showing inhibition of bacterial strains grown individually in the presence of each of the four strains of *Penicillium* sp. strain MB. Data for (a) *Staphylococcus*, (b) *Brevibacterium*, (c) *Brachybacterium*, and (d) *Psychrobacter* are shown. Points and connecting lines with standard errors of the means bars were used from two independent experiments with five biological replications each. In experiment 2, replication 5 was removed due to no growth of the bacteria in the control. Two-way analysis of variance (ANOVA) was performed for each bacterial community. Dunnett’s multiple comparison test was used and compared to the WT strain. (****) indicates a *p*-value <0.0001, (***) indicates a *p*-value <0.001, (**) indicates a *p*-value <0.01, (*) indicates a *p*-value <0.05, and no asterisk indicates not significant (ns). For exact *p*-values for each treatment, see Dataset 1.

*Penicillium* species often co-occur with yeasts in cheese rinds ^30, 39^ and the presence of another fungus may dampen the inhibitory effects of *Penicillium* sp. MB. To test this, we repeated all community experiments with the addition of the common cheese rind yeast *Debaryomyces hansenii*. We observed nearly identical patterns of inhibition of bacteria by MB_WT and loss of inhibition in MB_Δ*laeA* communities in the presence of *D. hansenii* (**Supplementary Fig. 3**).

To determine the role of LaeA in mediating pairwise interactions between *Penicillium* sp. MB and each of the individual bacterial species, we co-cultured each bacterium separately with the four *Penicillium* sp. MB strains on cheese curd agar. After 21 days, strains that had a functional LaeA (MB_WT, Hyg^R^, and MB_*laeA*c) inhibited the growth of all four bacteria (**Fig. 3)** with an inhibition hierarchy (log10 % decrease Alone vs. MB_WT) of *Psychrobacter* (100% inhibition) > *Brevibacterium* (73% inhibition) > *Staphylococcus* (24% inhibition) > *Brachybacterium* (21% inhibition). These data demonstrate that *Penicillium* sp. MB strongly and directly inhibits the growth of individual cheese rind bacteria, and this inhibitory effect is mediated by LaeA.

Collectively, these community and pairwise data demonstrate that LaeA regulates some aspect of *Penicillium* sp. MB physiology or metabolism that results in differential inhibition of bacterial species growth in the cheese rind community. The LaeA-regulated factor(s) have the ability to completely transform the composition of the bacterial community.

### RNA-sequencing reveals global alterations in expression of specialized metabolite genes in the Δ*laeA* strain

To identify SM biosynthetic gene clusters (BGCs) and other genes regulated by *laeA* in *Penicillium* sp. strain MB that might be driving bacterial-fungal interaction outcomes, we performed gene expression profiling using RNA sequencing (RNA-seq). RNA-seq libraries prepared from mycelium of MB_WT and MB_Δ*laeA* harvested 48 hours post-inoculation from CCA were examined for differential gene expression. This time point was selected because physiological differences between both strains were visually apparent (differences in pigmentation and spore production) and significant amounts and high-quality RNAs could be obtained. Using a log2 ratio cutoff of 2 (corrected *p*-value < 0.05), we identified 253 genes with decreased expression and 57 genes with increased expression in the MB_Δ*laeA* strain (**Fig. 4a**, **Dataset 2**). No transcripts for *laeA* were detected in the MB_Δ*laeA* strain confirming the successful deletion of this gene.

**Figure 4:**
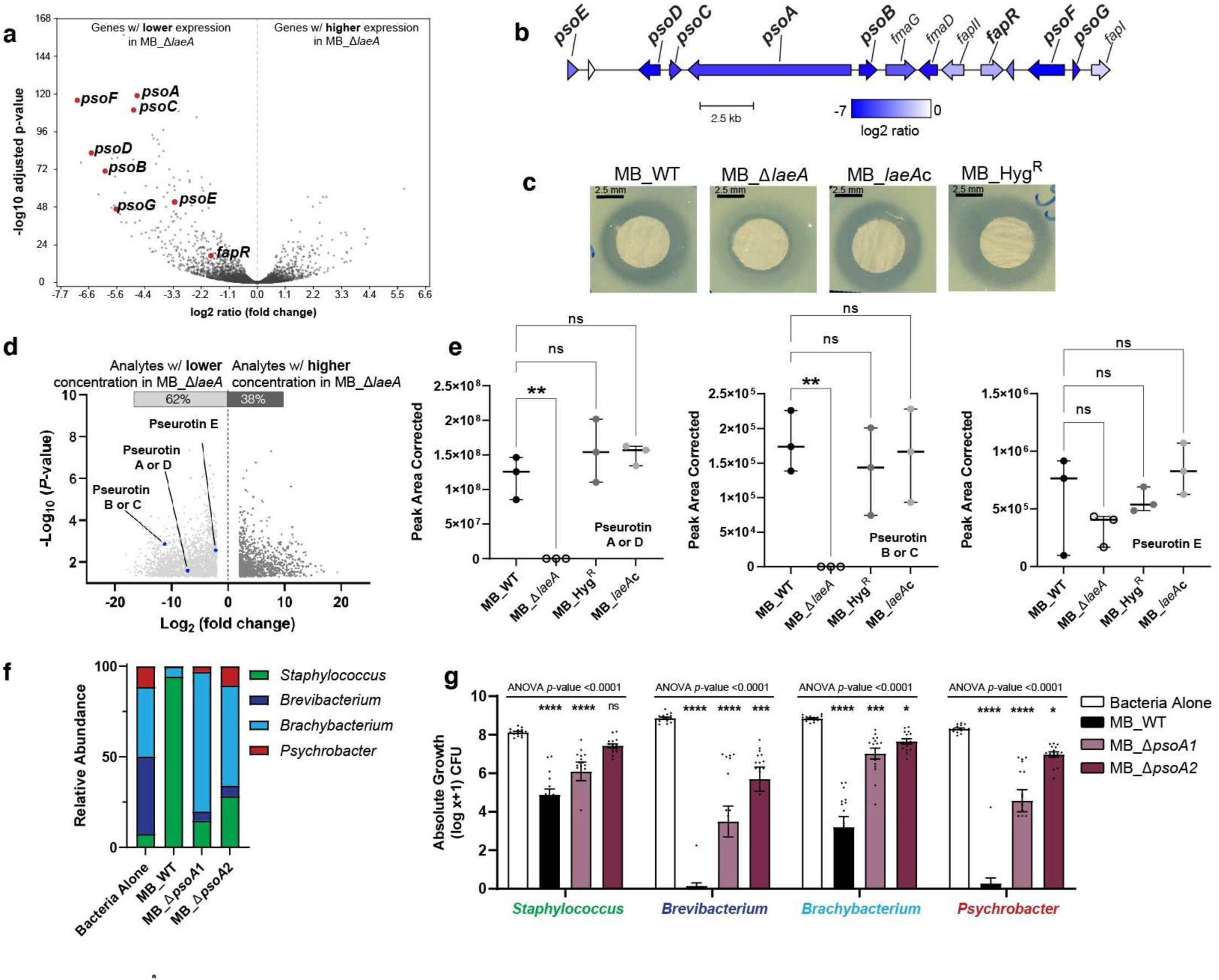
Deletion of *laeA* impairs production of metabolites with associated antimicrobial properties. **(a)** RNA-sequencing results showed a significant downregulation of biosynthetic genes in the pseurotin putative gene cluster when *laeA* was deleted. **(b)** Organization of the putative pseurotin biosynthetic gene cluster in *Penicillium* sp. strain MB and log2 ratio of expression in MB_Δ*laeA.* Gene names in bold are those known to be involved in pseurotin production in *Aspergillus* species. **(c)** Visualization of zones of inhibition of crude extracts collected from all four *Penicillium* isolates on YES medium against *Brachybacterium alimentarium* evaluated by the disk diffusion method. See **Supplementary Figure 7** for additional zone of inhibition data. **(d)** Volcano plots representing the number of analytes significantly regulated in MB_Δ*laeA* compared to the MB_WT strain on YES medium. The blue dots indicate pseurotin A and the other putative pseurotins identified in this analysis. **(e)** Comparison of the peak area corrected values for pseurotin A and the other putative pseurotins between strains. Each bar represents the means with +/- one standard error, and each dot represents a biological replicate (n = 3). One-way analysis of variance (ANOVA) was performed for each *Penicillium* strain. Dunnett’s multiple comparison test was used and compared to the MB_WT strain. (**) indicates a *p*-value < 0.01 and no asterisk indicates not significant (ns). **(f)** Inactivation of *psoA* led to loss of antibacterial activity and restored bacterial community composition. The compositions of MB_Δ*psoA*-1 and MB_Δ*psoA*-2 bacterial communities were not significantly different from one another, but were different from Bacteria Alone and MB_WT (PERMANOVA F = 49.97, *p* < 0.0001). (**g)** Histograms showing the inhibition of community members grown in the presence of the four strains of *Penicillium* sp. strain MB at 10 days post-inoculation. Data are mean relative abundance from three independent experiments with five biological replications each. In the box plots, the bar represents the standard errors of the means and each dot represents a biological replication. One-way analysis of variance (ANOVA) was performed for each bacterial community. Dunnett’s multiple comparison test was used and compared to the WT strain. (****) indicates a *p*-value <0.0001, (***) indicates a *p*-value < 0.001, (**) indicates a *p*-value < 0.01, (*) indicates a *p*-value < 0.05, and no asterisk indicates not significant (ns). For exact *p*-values for each treatment, see **Dataset 1**.

Using an enrichment analysis of GO terms in the list of differentially expressed genes, we identified depsipeptide, emericellamide, lactone, aspartic peptidase, indole alkaloid, fumagillin, epoxide, and alkaloid biosynthesis as pathways with greater than 30% of genes in a pathway being significantly downregulated (**Dataset 3**). All these biosynthetic pathways play major roles in specialized metabolite production of fungi. The GO terms glucose, hexose, monosaccharide, and nucleobase transporters were enriched in the upregulated genes of the MB_Δ*laeA* strain (**Dataset 3**).

We used the list of downregulated genes to identify specific BGCs that might be related to the observed inhibition of bacterial growth. BGCs were identified by locating groups of adjacent downregulated genes with annotations for hallmarks of biosynthetic gene clusters including polyketide synthases, nonribosomal peptide synthetases, terpene cyclases, and prenyltransferases^45^. The most downregulated BGC in the MB_Δ*laeA* strain contained genes homologous to the *A. fumigatus* fumagillin/pseurotin supercluster ^46^ (**Fig. 4a)**. Pseurotins have been reported as having antibacterial and insecticidal activities ^47^. The *Penicillium* sp. MB genome contains a 16-gene cluster with a complete set of predicted genes for pseurotin biosynthesis including *psoA*, *psoB*, *psoC*, *psoD*, *psoE*, *psoF*, and *psoG* (**Fig. 4b**). Some genes essential for fumagillin biosynthesis including the terpene cyclase gene (*fmaA*) are missing, suggesting the strain could synthesize pseurotin but not fumagillin (**Fig. 4b**). All these genes were highly downregulated (−4.7 to −7 log2 fold) and were among the most downregulated genes in the MB_Δ*laeA* strain (**Fig. 4a**). Other BGCs were downregulated in the MB_Δ*laeA* strain included a putative aspterric acid and quinolone BGC not known to have antibacterial activity as well as other BGCs encoding putative metabolites (**Dataset 2**). None of these BGCs were downregulated to the same extent as the pseurotin BGC.

### Loss of *laeA* alters the production of all members of the 1-oxa-7-aza-spiro[4,4]non-2-ene-4,6-dione class of antibacterial natural products

Our findings highlighted a significant role for *laeA* in the assembly of microbial communities that may be partly due to a fitness cost associated with physiological traits of this mutant (**Fig. 1**). The downregulation of several BGCs in the MB_Δ*laeA* strain suggested that differences in community structure between treatments could be due to antibacterial activities of *laeA*-regulated metabolites. To explore this hypothesis, we assessed metabolomic changes resulting from *laeA* deletion after 14 days of growth on four different media (CYA, PDA, YES and CCA). Crude extracts collected from all four *Penicillium* strains were later screened for antibacterial activity using the disk diffusion method against the same bacterial species used in the community and pairwise interaction assays.

Crude extracts collected from MB_WT growing on YES medium showed the highest antibacterial effects, with a clear circular zone of inhibition on all tested bacteria (**Fig 3c**; **Supplementary Fig. 4a**). Interestingly, crude extracts from MB_Δ*laeA* cultures generated significantly smaller zones of inhibition than extracts from control strains regardless of the bacterial strain tested (**Supplementary Fig. 4a**). Crude extracts collected from cultures of the MB_Δ*laeA* and the control strains did not show significant differences in their ability to inhibit bacterial growth, except against *Psychrobacter* (**Supplementary Fig. 4b-d)**. Extracts collected from CCA cultures showed the lowest inhibitory effects on all tested bacteria, likely due to the high amount of fat in this medium resulting in low recovery of metabolites from the organic phase (**Supplementary Fig. 4d**). Similar problems with metabolite recovery have previously been attributed to the high fat content in milk. ^48^ Therefore, we focused on exploring the metabolomic changes on YES medium to pinpoint the *laeA*-regulated metabolite(s) responsible for the shift in bacterial community composition.

Analysis of metabolomic data on YES medium showed that the deletion of *laeA* led to a 2-fold decrease in the production of 62% of the 2703 significantly differentially produced analytes and 2-fold increase in the production of of 38% of all significantly differentially produced analytes in negative ionization mode (**Fig. 4d**). Pseurotins have a 1-oxa-7-aza-spiro[4,4]non-2-ene-4,6-dione core and multiple forms exist (A through F) with slight modifications to this core ^49^. Using a list of chemical formulas assigned to known specialized metabolites produced by this *Penicillium* species and closely related species, we were able to identify peaks that correspond to putative pseurotins. Strikingly, the antibacterial metabolite pseurotin A showed a 140-fold decrease in the Δ*laeA* mutant compared to the WT strain (*p*-value ∼ 0.02) (**Fig. 4e**). The identification of pseurotin A (C_22_H_25_NO_8_) was confirmed by High-resolution mass spectrometry (HRMS) [M-H]^-^ (m/z 430.1503, calculated for C_22_H_24_NO_8_ 430.1507) and by HRMS/MS fragmentation. The mass spectrum showed the two fragmentation ions (m/z 270.0768 and 308.1137) consistent with the mass spectrum ion previously reported for pseurotin A^46^. We could not perform experimental confirmation of other putative pseurotins due to the lack of commercially available standards or fragmentation databases. The previously published chemical formulas of pseurotin B, C, D and E^50^ were applied to MAVEN, which identified a significant reduction of production of these putative metabolites in the absence of *laeA* (**Fig. 4e**).

The significant decrease in the synthesis of pseurotin A and the other pathway derivatives on YES medium matched the transcriptomic data showing a downregulation of the pseurotin gene cluster in the MB_Δ*laeA* strain (**Fig. 4a**). Combined with previous reports of antibacterial properties of pseurotin,^47^ our concurring datasets suggested that pseurotins could explain the striking antibacterial activity of this *Penicillium* isolate.

### Inactivation of pseurotin production leads to a loss of antibacterial activity and restores bacterial community composition

To test whether pseurotins were the fungal metabolites regulating bacterial community structure, we disrupted the hybrid PKS-NRPS synthase gene *psoA* required for pseurotin production in *Penicillium* sp. strain MB using a CRISPR-Cas9 system (**Supplementary Fig. 5b**). The disruption of this gene in two independent mutants (MB_Δ*psoA*-1 and MB_Δ*psoA*-2) and the absence of Cas9 off-target cleavage events were confirmed using whole-genome resequencing (**Supplemental Fig. 5c**). LC-MS/MS analysis of MB_Δ*psoA*-1 and MB_Δ*psoA*-2 confirmed a lack of pseurotin A production (**Supplementary Fig. 6**). In contrast, MB_WT produced on average 1.359 mg of pseurotin A (**Dataset 4**).

When we repeated the bacterial community assembly assays described above with the Δ*psoA* mutants, we saw two striking patterns. First, the two mutants did not reduce total bacterial growth as much as the MB_WT strain; while MB_WT had a 34.7% reduction in total CFUs across all four bacterial species, the average total bacterial reduction across the two pseurotin mutants was only 13.9% (**Fig. 4g**). As we observed earlier (**Fig 2**), *Brevibacterium*, *Brachybacterium*, and *Psychrobacter* were most inhibited by the MB_WT strain. Similar to co-culture with MB_Δ*laeA*, the Δ*psoA* mutants had minor inhibitory impacts on these bacterial genera (**Fig. 4g**). Second, we observed that bacterial community composition was significantly different between the MB_WT and the MB_Δ*psoA*-1 and MB_Δ*psoA*-2 mutant communities. As with the Δ*laeA* knockout above, the bacterial community composition shifted from dominance of *Staphylococcus* in the MB_WT community to a mix of all bacterial species in the MB_Δ*psoA*-1 and MB_Δ*psoA*-2 communities (**Fig. 4f**). These data combined with our transcriptomic and metabolomic data above demonstrate that the pseurotin BGC is regulated by LaeA in this *Penicillium* species and that the loss of pseurotin production eliminates much of the strong inhibitory effect of this fungus on bacterial community assembly.

### Components of the pseurotin biosynthetic gene cluster are widely distributed across Ascomycota

Considering the strong ecological impacts of the pseurotin BGC, we used comparative genomics to identify other fungi whose genomes encode this cluster or closely related clusters with similar antimicrobial properties. Previous studies have demonstrated that the pseurotin/fumagillin supercluster was subject to rearrangements and gene loss in *A. clavatus*, *Neosartorya fischeri,* and *Metarhizium anisopliae*^46^. Scanning across a kingdom-level sample of fungal genomes, we found that pseurotin genes are only present in the Ascomycota (**Fig. 5; Supplementary Fig. 7**). Only a small subset of fungi are predicted to have all eight pseurotin genes, including species in the genera *Aspergillus*, *Penicillium*, *Metarhizium*, *Tolypocladium*, and *Venustampulla* (**Dataset 5**). Of the 77 species of *Aspergillus* in this dataset, only 11 have all 8 genes. Of the 24 species of *Penicillium*, only 2 (in addition to *Penicillium* sp. MB) have all 8 genes (**Fig. 5a**). Another closely related *Penicillium* strain that we isolated from a separate cheese production facility in the USA (*Penicillium* sp. 261) also has the full pseurotin BGC (**Fig. 5b**).

**Figure 5.**
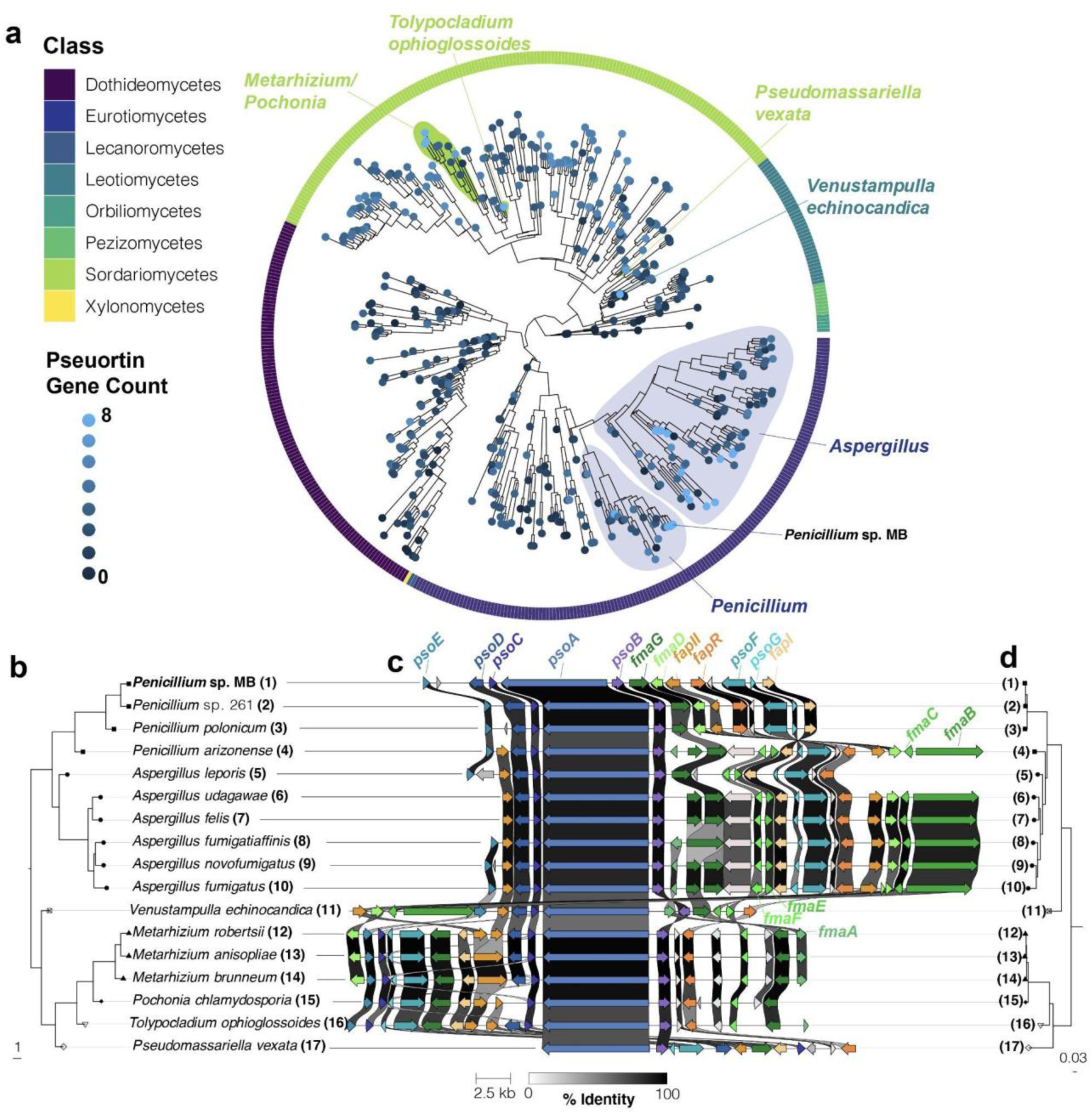
Components of the pseurotin biosynthetic gene cluster are widespread across Ascomycota, but the full cluster is mainly found in *Aspergillus* and *Penicillium* species. **(a)** The subclade of Ascomycota identified as having higher prevalence of pseurotin cluster genes (**Supplementary Figure 7**) presented with a single leaf per species. Phylogenetic relationships were determined from the consensus of 290 maximum-likelihood trees constructed for benchmarking single copy orthologs (BUSCOs). The outer ring indicates the taxonomic order that species pertain to. Terminal branch lengths are not calculated. The number of pseurotin genes was determined using reciprocal best-hit BLAST analysis using the *Aspergillus fumigatus* pseurotin genes *psoA, psoB, psoC, psoD, psoE, psoF, psoG*, and *fapR* and is indicated as the intensity of the tip color. **(b)** A phylogeny constructed in the same way as (a) representing a subset of species identified as having seven or eight pseurotin genes that were manually selected to represent taxonomic breadth. **(c)** An alignment of pseurotin gene clusters. Genes in the pseurotin cluster are labeled in blue while constituents of the intertwined fumagillin cluster are indicated in green. Regulatory genes of these clusters are labeled in orange. Coloration of genes is based on homology searches and visualizations performed by clinker. The weight of lines between homologous genes indicates the percent identity; genes sharing identity below 30% are not indicated. **(d)** A maximum likelihood phylogeny constructed from the concatenation of PsoA, PsoB, and PsoC sequences. Note that phylogenetic relationships between (d) and (b) are compatible, suggesting vertical transmission of the pseurotin gene cluster. The strain information of the species used in this analysis are provided in **Supplementary Table 1.**

Given the taxonomic breadth of fungi containing the pseurotin cluster, we sought to understand the evolutionary origins of this cluster within and outside the Ascomycota, including whether horizontal or vertical transmission explains patterns of pseurotin gene distribution. The overall history of this BGC appears consistent with vertical transmission and gene loss in some lineages; the topology of a phylogeny constructed from 290 benchmarking single copy orthologs (BUSCOs) was compatible with another phylogeny constructed using PsoA, PsoB, and PsoC sequences (**Fig. 5b-d**). We cannot entirely rule out the possibility of horizontal gene transfer, but such transfers would have to be ancient to allow for BGC and species histories to align. Patterns of vertical transmission are also apparent in the synteny of the BGC. For example, an inversion of several genes at one end of the cluster (*fmaD-fapI*) that is present in most *Penicillium* spp. but not in *Aspergillus* spp. is missing in the early-divergent *Penicillium arizonense* (**Fig. 5c**), suggesting that this mutation occurred sometime after the diversification of this genus.

To explore the possibility of an ancient horizontal gene transfer event from outside the fungi, we BLASTed the PsoA protein against all NCBI databases excluding fungi. The highest-scoring hit from this search (94% of the query cover, 32.38% identity) was an uncharacterized hypothetical protein that is annotated as PKS in the plant *Carpinus fangiana* (accession KAB8336832.1). However, the reciprocal BLAST of this protein against fungi found better hits (100% coverage and ∼60% identity) against other uncharacterized PKSes and putative lovastatin nonaketide synthases. While our results demonstrate that the pseurotin BGC is widespread in a subclade of Ascomycota, further analyses exploring the evolution of this gene cluster are needed.

## DISCUSSION

The chemistry, genetics and physiological effects of fungal specialized metabolism are widely studied and recognized, but their ecological consequences in microbiomes are largely unknown. Previous genetic and -*omics* approaches identified key metabolites and pathways regulated by LaeA ^28, 51–54^, but these data have not been integrated into a comprehensive model for studying microbial interactions that influence community assembly. By toggling LaeA between on and off, we were able to identify one class of metabolites produced by a strongly inhibitory fungus that allows it to dramatically remodel the bacterial communities of cheese. Our integration of physiological assays, transcriptomics, and metabolomics, more broadly demonstrates how LaeA can be used to identify fungal metabolites that control the assembly of microbiomes. Other LaeA-regulated metabolites and fungal traits likely impact bacterial community assembly in this system, but it was striking how much of the bacterial community was determined by the function of a single BGC.

The intertwined pseurotin and fumagillin supercluster was initially identified in *Aspergillus fumigatus* and reported to be under the control of LaeA^46^. Our study shows that this BGC exists in the cheese isolated *Penicillium* strain MB and is also positively regulated by LaeA **(Fig. 4a, b).** This metabolite was previously found to exhibit a range of bioactivities at moderate to high concentrations (up to 50 µg/mL) with potential therapeutic applications due to its immunosuppressive activity^55^, antibacterial activity ^47, 56^, nematocidal activity ^57^, insecticidal activity^58^, antiparasitic as well as anticancer activity ^59^. Previous studies of antibacterial activity have focused on purified versions of the compound and have not used natural production from a host for assays. In our analysis, the production of pseurotin A by the MB_WT strain was roughly 2.7 times higher than the concentration used in the aforementioned studies (**Dataset 4**) and may explain why this fungus has such strong inhibitory effects on bacterial communities. Characterization of how other fungi that produce pseurotins impact bacterial community assembly will further clarify how this class of metabolites plays roles in multispecies microbiomes.

In addition to its regulation and role in the cheese *Penicillium* isolate, we have found that components of the pseurotin BGC can be found intermittently in the Ascomycota, but not in any other fungal phyla. Pseurotin A has been reported in several different filamentous fungi including species of *Aspergillus*, *Penicillium,* and more recently the distantly related taxon *Metarhizium* ^58, 60–62^. Our finding of all eight genes of this BGC in several other genera (*Tolypocladium* and *Venustampulla*) suggests that fungal pseurotins may be widely distributed across a range of fungal niches, from food production and indoor environments to forest soils. Additional studies of these fungi and neighboring bacteria are needed to fully characterize the ecological significance of pseurotins across ecosystems.

Every time a consumer eats a piece of a naturally aged cheese, they ingest the many metabolites secreted into the cheese by rind microbes. Surprisingly little is known about the diversity and functions of these microbial metabolites ^37, 63–66^. Our work demonstrates that fungal SMs produced in cheese environments can serve as mediators of microbiome formation at concentrations relevant to naturally aged cheeses. The interactions described here occurred in a highly controlled laboratory environment and their relevance in full-scale cheese production remains to be determined. The *Penicillium* sp. strain MB used in this study was causing cheese production problems by inhibiting normal rind formation, suggesting that the disrupted community assembly observed in the lab could also happen in cheese caves. Many other filamentous fungi found in cheese rinds possess uncharacterized secondary metabolites that might be responsible for similar negative outcomes. Systematic exploration of LaeA-regulated metabolites and their ecological roles will not only discover microbial mechanisms underlying traditional cheese-making but will also illuminate how fungal SMs mediate microbiome composition in all environmental niches where fungi reside.

## METHODS

### Microbial strain isolation and culture conditions

The *Penicillium* sp. strain MB used in this study was isolated from a natural rind cheese made in the United States. Its genome has been deposited in NCBI with accession number GCA_008931935.1. Previous comparative genomic analyses demonstrated that this strain is in the *Penicillium* section *Fasciculata*, which includes *Penicillium camemberti* and other food-associated species ^42^. Preliminary observations demonstrated that this *Penicillium* strain had potent antibacterial activity and was therefore an interesting strain to explore microbial interactions and secondary metabolite regulation through the inactivation of the global regulator of fungal secondary metabolism, LaeA.

To prepare a fresh spore suspension prior to experiments, the *Penicillium* strain MB was activated on glucose minimal media (GMM) ^67^ for 7 days at 25 °C and spores were harvested in 0.01% Tween 80 and counted using a hemocytometer.

Four bacterial strains (*Staphylococcus equorum* strain BC9*, Brevibacterium linens* strain JB5*, Brachybacterium alimentarium* strain JB7, and *Psychrobacter sp*. strain JB193) were used in antibacterial assays. These species span three bacterial phyla (Firmicutes, Actinobacteria, and Proteobacteria) ^39^ that are most abundant in cheese rinds. They have been used as a model community in previous work from our lab ^40, 41, 68^ and have been demonstrated to have varying responses to the presence of *Penicillium* ^39, 41^. This model community also has a well-defined community succession with *Staphylococcus* and *Psychrobacter* dominating early in succession (days 0-10) and the Actinobacteria *Brevibacterium* and *Brachybacterium* dominating later in succession (days 10-21). Bacterial strains were also maintained as glycerol stocks and streaked on brain heart infusion (BHI) agar prior to experiments.

### Construction of gene deletion cassettes

To knockout *laeA* in the *Penicillium sp.* strain MB, the isolate was first subjected to antimicrobial susceptibility testing towards hygromycin and phleomycin, two antibiotics commonly used by our group as selectable markers in fungal transformations. The isolate showed a confirmed sensitivity to both antibiotics. A three round PCR deletion strategy was used to replace the *laeA* ORF in the *Penicillium sp.* strain MB with the *hph* gene, whose expression confers selection on hygromycin. The schematic representation of the *laeA* gene replacement with the *hph* gene is depicted in **Supplementary Fig. 1a**. Each deletion cassette (5’flank-*hph*- 3’flank) was constructed using three sequential PCR reactions. In the first PCR round, about 1kb of genomic sequence that flanks either the 5’ or 3’ end of the *laeA* ORF was amplified from strain MB using respectively the primer set PMB_KOlaeA_5’ or 3’F/R. The *hph* gene was amplified from plasmid pUCH2-8 using primers hph_F and hph_R. A second PCR reaction was performed to assemble by homologous recombination the three individual fragments from the first round PCR. The deletion cassettes were finally amplified using nested primer sets (PMB_KOlaeA_NestedF/R).

To test whether pseurotin was involved in the antimicrobial activity and shifts in bacterial community composition observed with this *Penicillium* strain, we knocked out the *psoA* gene that encodes for the hybrid polyketide synthase-non-ribosomal peptide synthetase (PKS/NRPS) of the putative pseurotin biosynthetic gene cluster to disrupt the production and accumulation of all pseurotin. The same strategy described earlier was adopted for the construction of the *psoA* deletion cassette. The primer sets PMB_KOpsoA_5’/R and 3’F/R were used to amplify the 1kb homology arms flanking the *psoA* gene on the 5’ and 3’ ends, respectively. The *hph* gene under the control of the Tef1 promoter and terminator was amplified from the plasmid pCS01 using the primer set Hyg-tef1F/R. The plasmid map is given and annotated in **Supplementary Fig. 5a** second PCR reaction was performed to assemble by homologous recombination the three individual fragments from the first round PCR. The deletion cassette was finally amplified using the nested primer set (PMB_psoA_nestedF/R). The sequences of the primer sets used for the construction of the deletion cassettes are shown in **Supplementary Table 2.**

### *PEG-*mediated protoplast transformation

To generate the deletion strains, a protoplast-mediated transformation protocol routinely used by our group was employed and optimized to achieve a successful protoplasting of this *Penicillium sp.* strain. Briefly, 10^9^ fresh spores from each strain are cultured in 500 mL of GMM broth supplemented with 1g/L yeast extract for 12 h under 25 °C and 280 rpm. Newly born hyphae were harvested by centrifugation at 8000 rpm for 15 min and hydrolyzed in a mixture of 30 mg lysing enzyme from *Trichoderma harzianum* (Sigma-Aldrich) and 20 mg Yatalase (Fisher Scientific) in 10 mL of osmotic medium (1.2 M MgSO_4_ and 10 mM sodium phosphate buffer). The quality of protoplast was monitored under the microscope after four hours of shaking at 28 °C and 80 rpm. The protoplast mixture was later overlaid gently with 10 mL of chilled trapping buffer (0.6 M sorbitol and 0.1 M Tris-HCl, pH 7.0) and centrifuged for 15 min under 4°C and 5000 rpm. Protoplasts were collected from the interface, overlaid with an equal volume of chilled STC (1M sorbitol, 10 mM Tris-HCL and 10 mM CaCl_2_) and decanted by centrifugation at 6000 rpm for 10 min. The protoplast pellet was resuspended in 500 µL STC and used for transformation. For protoplast transfection, 100 μL of freshly isolated protoplasts and 5 μg of linear DNA containing the deletion cassette were mixed to a final volume of 200 µl of STC buffer. The contents were mixed by gently inverting the tubes. After 50 min incubation on ice, 1.2 ml of 60% (w/v) PEG solution (60 g PEG 3350, 50 mM CaCl_2_ and 50 mM Tris-HCl, pH 7.5) were added to the mixture and incubated for an additional 20 min at room temperature. The mixture was supplemented with 5ml of STC and mixed into 50 ml of SMM top agar (GMM supplemented with 1.2 M sorbitol) containing hygromycin at a final concentration of 150 µg/ml. The mixture was inverted several times and each 5 ml were poured onto a selective SMM bottom agar plate. The transformation plates were incubated at 25°C for 5-7 days.

### CRISPR/Cas9-mediated knockout of psoA gene

The large size of the *psoA* gene (∼12 kb) made the standard transformation method described for *laeA* deletion ineffective. Therefore, we switched to a ribonucleoprotein-CRISPR-Cas9 (RNP-CRISPR-Cas9) system. The schematic representation of the CRISPR cas9 system used for engineering the *psoA* knockout strain is depicted in **Supplementary Fig. 5b**. The fungal transformation steps followed were the same as described above, the only difference being the co-delivery of the Cas9-gRNA RNP complex along with the linear donor DNA (*psoA* deletion cassette) to the protoplasts. The Crispr RNA (crRNA) was designed on the PKS-NRPS conserved domain in the *psoA* sequence using the CRISPOR webtool (http://crispor.tefor.net/) that offers off-target and efficiency predictions. The selected crRNA (5’ GGAUCGAUCUUGAACAGCAG 3’) showed a predicted targeting efficiency of 100% and 0 predicted off targets. This crRNA was subjected to an additional confirmation using the RNAfold web server (http://rna.tbi.univie.ac.at/cgi-bin/RNAWebSuite/RNAfold.cgi) that allows to determine the gRNA secondary structure. The interpretation of the results and validation of the gRNA were done following the guidance by Hassan et al. ^69^. The crRNA and tracrRNA (IDT, cat no. 1072534) were synthesized using the Alt-R CRISPR-Cas9 system from Integrated DNA Technologies (IDT) (San Diego, California). The crRNA and tracrRNA were mixed at a concentration of 200 nM each in IDT duplex buffer in a final volume of 20 µL. The gRNA complex was formed by incubation at 95 °C for 5 min, followed by slowly cooling down for 20 minutes at room temperature, and then stored at −20 °C. To form the RNP complex, 8.3 µL of HiFi Cas9 Nuclease (IDT, cat no.1081060) and 6 µL of the previously made gRNA were mixed in nuclease free water to 25 μL final volume. The mixture was incubated for 15 min at room temperature prior to use.

### Confirmation of gene-deletion strains

After 5-7 days of incubation at 25°C, colonies grown on SMM plates supplemented with hygromycin (150 µg/ml) were subjected to a second round of selection on hygromycin. Single-spored transformants were later tested for proper homologous recombination at the ORF locus by PCR.

The correct replacement of the *laeA* with the *hph* gene was first verified by PCR analysis of genomic DNA derived from the transformant strains using primers that amplify the *laeA* ORF. One ORF-specific confirmation primer set (PMB_laeA_F/R) was chosen for the strain. The positive deletion strains identified by PCR were further checked for a single insertion of the deletion cassette by southern blot analysis. Probes corresponding to the 5’ and 3’ flanks of the *laeA* gene in each strain were labeled using [α32P] dCTP (PerkinElmer, USA) following the manufacturer’s instructions.

The confirmation of the Δ*psoA* mutants was first done by screening for the absence of the ORF using the primer set PMB_psoA_F/R. Later, identification of exact gene deletion locations and assessment of off-target effects of the cas9 were analyzed by whole-genome re-sequencing of the PCR positive knockout strains. DNA was extracted from each Δ*psoA* strain using a Qiagen DNAEasy PowerSoil Extraction kit. DNA was sent to the Microbial Genome Sequencing Center for library preparation and sequencing on an Illumina NextSeq 2000. To determine the specific location of CRISPR deletion, reads were mapped to the *Penicillium* sp. MB_WT genome using Bowtie. To assess whether any off-target mutations were caused by the CRISPR deletion of *psoA*, FreeBayes was used to identify variants in the mappings of both Δ*psoA* strains.

### Construction and confirmation of complement strains

To confirm that the phenotype exhibited by the MB Δ*laeA* strain is caused by the deletion of this specific gene, the MB Δ*laeA* strain was complemented with a WT copy of the *laeA* gene using phleomycin as a selectable marker. Restriction sites for NotI were introduced at the predicted native promoter and terminator of the *laeA* gene using primers PMB_laeAcomp_F and PMB_laeAcomp_R. The *laeA* gene was cloned into the pBC-phleo plasmid at the multiple cloning site located within the *lacZ* gene (**Supplementary Fig. 1c**). The ligation of the digested insert into the recipient plasmid was performed using T4 DNA ligase (New England Biolabs) following the manufacturer’s instructions. The ligation reaction was later transformed into *Escherichia coli* DH5alpha competent cells following the manufacturer’s directions (Thermo Fisher Scientific). Five white bacterial colonies were randomly selected from the blue-white screening LB agar plate and screened for successful ligations by conducting a diagnostic restriction digest with the NotI restriction enzyme. The *E. coli* strain carrying the correct plasmid (labelled PJT3) was then grown in 50 mL Lysogeny broth (LB) supplemented with chloramphenicol (35 μg/mL), and the plasmid DNA was isolated using Quantum Prep®Plasmid Midiprep Kit (Biorad) according to the manufacturer’s’ instructions. Ten micrograms of plasmid DNA were used for the transformation of the MB_Δ*laeA* strain following the same protocol described above. Prior to transformation, the plasmid was linearized using the NotI restriction enzyme.

Southern blotting was performed to confirm the single integration of *laeA* into the MB deletion strain using the *laeA* ORF sequence (amplified using the primer set laeA_ORF_F/R) for making the probe. Genomic DNAs from both *laeA* complement and deletion strains were digested with the same enzyme used for cloning. A positive control corresponding to the PJT3 plasmid was incorporated in the southern blot analysis, and the MB_Δ*laeA* strain was used as a negative control.

### Morpho-physiological analysis

The impact of *laeA* deletion on the morpho-physiological traits of the cheese *Penicillium* sp. MB was evaluated by monitoring the growth, sporulation, and germination of the MB_WT strain in comparison to the MB_Δ*laeA,* MB_*laeA*c and the MB_Hyg^R^ control strains. The phenotypic appearance and vegetative growth were evaluated on 5 different media, which are: GMM (Glucose Minimal Medium), CYA (Czapek Yeast Extract Agar), YES (Yeast Extract Sucrose Agar), PDA (Potato Dextrose Agar) and CCA (Cheese Curd Agar). The composition of all the media used is provided in **Supplementary Note 1.**

Spore production and germination were assessed on GMM agar and broth, respectively. For conidial counts, fresh spores from each strain were diluted to 10^5^ spores per ml in GMM top agar and overlaid onto agar plates of the same medium. The plates were incubated at 25 °C and agar plugs removed on the second, third- and fourth-day post inoculation were homogenized in 3 mL of 0.01% Tween 80 using the VWR 200 homogenizer. Total spore counts were made using a hemocytometer. Conidial germination rates were evaluated over a 24-hour growth period using a Nikon Ti inverted microscope. A spore suspension of 10^5^ spores per ml of GMM broth was prepared for each strain and about 1mL was distributed into 3 replicate wells of a 24-sterile well plate. Five pictures per well were taken an hour apart beginning 4 hours post-incubation. The number of germlings were counted for each strain and the percent of germinated spores was plotted against time to estimate the germination rates.

### Community and pairwise interaction assays

To determine how the deletion of *laeA* impacted microbial community assembly, we reconstructed cheese rind bacterial communities on cheese curd agar (CCA) with each of the *Penicillium* strains and measured total bacterial community abundance (as total CFUs) and bacterial community composition (relative abundance of each community member) at 3, 10 and 21 days of community assembly. Each member of a four-member bacterial community (*Staphylococcus equorum* BC9*, Brevibacterium auranticum* JB5, *Brachybacterium alimentarium* JB7*, and Psychrobacter sp.* JB193) was initially inoculated at 200 CFUs per species in five treatments: **1)** bacteria alone, **2)** bacteria + *Penicillium* WT strain, **3)** bacteria + *Penicillium* Δ*laeA* strain (MB_Δ*laeA)*, **4)** bacteria + *Penicillium* hygromycin resistance control strain (MB_Hyg^R^), and **5)** bacteria + *Penicillium laeA* complement strain (MB_*laeA*c)*. Penicillium* strains were also inoculated at 200 CFUs from experimental glycerol stocks ^43^. For each treatment, replicate communities were inoculated on the surface of 150 µL of cheese curd agar dispensed into each well of a 96-well plate. To determine bacterial community composition, communities were harvested from individual wells with a sterile toothpick, suspended in 500 µL of phosphate buffered saline in a 1.5 mL microcentrifuge tube, homogenized with a sterile micropestle, and serially diluted onto plate count agar with milk and salt (PCAMS) media ^43^. To selectively plate bacteria, 100 mg/L of cycloheximide was added to PCAMS. To quantify *Penicillium* abundance, 50 mg/L of chloramphenicol was added to PCAMS. Each of the four bacteria have very distinct colony morphologies making it easy to determine the abundance of each community member.

Pairwise interactions between each individual bacterium and the four *Penicillium* strains (MB_WT, MB_Δ*laeA*, MB_Hyg^R^, and MB_*laeA*c) were assessed using the same experimental setup as the community experiments. Each bacterium was inoculated on the surface of a well of a 96-well plate with PCAMS either alone or with 200 CFUs of each of the four *Penicillium* strains. Bacterial abundance was determined at 3, 10 and 21 days by plating harvested co-cultures on PCAMS supplement with 100 mg/L of cycloheximide.

To determine the role of pseurotin in shaping cheese microbial communities, these community assays were conducted with the Δ*psoA*-1 and Δ*psoA*-2 strains. The experimental setup and data collection and analysis were identical to the experiments with the MB_WT, MB_Δ*laeA*, MB_Hyg^R^, and MB_*laeAc* strains noted above.

### RNA-sequencing analysis

Transcriptome changes in *Penicillium* sp. strains MB_WT and MB_Δ*laeA* were investigated using RNA-sequencing analysis of cultures growing on CCA medium. Inoculum of both strains were prepared from 1-week cultures on PCAMS medium. A 1 cm^2^ plug was taken from the leading edge of mycelium and homogenized in 500 µL of phosphate buffered saline (PBS). A 20 µl inoculum was spotted onto a CCA plate at three evenly spaced positions. After 48 hours of growth in the dark at 24 °C, the spots were about 1.5 cm in width. The MB_WT had produced blue colored spores whereas the MB_Δ*laeA* spores were lighter in color. The entire fungal growth from each spot was cut-off from the CCA plates, placed in RNAlater (Qiagen) and stored at −80 °C. Four biological replicates were sampled for each strain.

RNA was extracted from one of the three spots from each replicate plate using the Qiagen RNeasy Plant Mini Kit after grinding the sample in liquid nitrogen. Approximately 100 mg of ground fungal biomass was mixed in 750μl of Buffer RLT supplemented with 10μl of β-mercaptoethanol. The manufacturer’s recommended protocol was followed for RNA extraction, including an on-column DNAse treatment. To isolate mRNA, the NEBNext ® Poly (A) mRNA Magnetic Isolation Module (New England Biolabs) was used. This mRNA was used to generate RNA-seq libraries using the NEBNext ® Ultra II RNA Library Prep Kit for Illumina following the manufacturer’s recommended protocol. The RNA-seq libraries were sequenced using 125 bp paired-end Illumina sequencing on a HiSeq at the Harvard Bauer Core.

Duplicate reads were removed, and the total number of reads was subsampled to 3.8 million forward reads that were used for read mapping and differential expression analysis. Reads were mapped to a draft genome of *Penicillium* sp. strain MB. Read mapping was performed with TopHat v2.1.0 ^70^. Differentially expressed genes were identified using DeSeq2 ^71^. Genes with greater than 5-fold change in expression and FDR corrected *p*-values of < 0.05 were considered as differentially expressed. To identify specific biological pathways that were enriched in the sets of downregulated or upregulated genes, we used a KOBAS 2.0 ^72^ to conduct a hypergeometric test on functional assignments from the gene ontology (GO) database (using the *Aspergillus flavus* genome as a reference for GO ID assignment) with Benjamini and Hochberg FDR correction.

### Metabolite profiling by UHPLC-MS analysis

To determine the effect of *laeA* deletion on the biosynthetic metabolome of the *Penicillium* sp. strain MB, all four strains (MB_WT, MB_Δ*laeA*, MB_Hyg^R^, and MB_*laeAc*) were cultivated by centrally inoculating 10^6^ fresh spores on 60 mm petri dishes containing 10 ml of the agar media PDA, CYA, YES and CCA. Three technical replicates per strain and condition were prepared. The cultures were incubated at 25 °C for 2 weeks. After the incubation period, all cultures were freeze-dried (approximately 3 g dry weight) and grinded into 5 mL sterile water. Soluble metabolites were later extracted by solvent extraction procedure using 5 mL of ethyl acetate. An organic metabolite fraction was generated by liquid:liquid partitioning and dried under vacuum. The crude extract was then dissolved in 400 µL acetonitrile:water (80:20 v/v) at a concentration of 100 µg/µl. Samples were later analyzed by UHPLC-MS as previously described ^73^. The total dataset was first evaluated using the software MAVEN and the XCMS open-source package. Differential masses found via XCMS were filtered by having a maximum intensity greater than 4×10^4^. Identified masses that had a maximum intensity lower than 4×10^4^ were considered as a background. The volcano plot was later constructed to determine statistically significant data points in crude extracts analyzed for both MB_WT and MB_Δ*laeA* in negative ionization mode. For volcano plot construction, metabolites were filtered based on a *p*-value lower than 0.05 and a fold change higher than 2 and lower than −2.

To confirm the lack of pseurotin production by the two Δ*psoA* mutants, crude extracts from 14-day cultures on YES agar medium of both MB_ Δ*psoA* and MB_WT were obtained and assessed by high-resolution parallel reaction monitoring (PRM) LC-MS/MS using a Vanquish uHPLC plumbed directly to a Q Exactive Plus mass spectrometer (Thermo Scientific) outfitted with a 75 µm ID nanospray emitter packed with 15 cm Kinetex C18 resin (1.7 µm particle size; Phenomenex). Mobile phases included solvent A (95% H_2_O, 5% acetonitrile, 0.1% formic acid) and solvent B (30% H_2_O, 70% acetonitrile, 0.1% formic acid). Nanoliter flow rates were achieved by uHPLC split-flow and measured 300 nL/min at the nanospray emitter. Five microliters of each sample was auto-injected prior to the split, leading to the separation and analysis of 10 nL of extract over a 30 min chromatographic method: 0 to 100% B over 17 min; 100 to 0% B over 3 min; hold at 0% B for 10 min to re-equilibrate the column. Pseurotin A was targeted for PRM analysis in positive ion mode with a duty cycle that included a selected ion monitoring scan (427 - 437 m/z range; resolution 35,000; 2 microscan spectrum averaging) followed by a PRM scan targeting the pseurotin A ion (m/z of 432.1653 [M+H]^+^; resolution 17,500; 2.0 m/z isolation window with a 0.5 m/z offset; normalized collision energy [NCE] in the HCD cell at 35). Sample extracts were measured across three technical replicates of each strain (WT_MB, MB_Δ*psoA1,* MB_Δ*psoA2)*, including 10 µM pseurotin A standard (Cayman Chemical). Standard addition experiments were also performed using the above PRM method on samples extracted from YES grown MB_WT strain to assess the pseurotin A concentration. All PRM data was analyzed using Skyline ^74^ and FreeStyle (Thermo Scientific) software.

### *In vitro* antimicrobial assay

To determine whether the findings observed with community and pairwise interaction assays are due to secreted metabolite(s), the antimicrobial activities of all crude exudates collected from cultures of MB_WT, MB_Δ*laeA*, MB_hyg^R^ and MB_*laeA*c strains on various media were evaluated using the paper disk agar diffusion method. The antimicrobial properties were assessed against the same bacterial strains used for community and pairwise interaction assays except the *B. linens* strain JB5 due to the inability of growing this strain for these *in vitro* experiments. Bacterial strains were first cultured in 5 ml of LB broth under 280 rpm at room temperature for 24 - 48 hours. The optical density of the bacterial suspension was later adjusted to 1. One milliliter of the bacterial suspension was then added to 20 mL of LB top agar and 5 mL were gently applied on agar dishes of the same medium. In sequence, sterile disks impregnated with 10 µL of extracts (at a concentration of 100 µg/µl) dissolved in acetonitrile:water (80:20 v/v) were placed over the bacterial culture plates. One disk containing the solvent previously used for resuspension was used as negative control. For each bacterium, one disk of ampicillin at a concentration of 100 µg/mL was applied as a positive control. All dishes were incubated at room temperature, for 24 - 48 hours. At the end of the incubation period, each dish was examined, and inhibition halo diameters were measured.

### Whole kingdom analysis of pseurotin gene cluster

To contextualize the ecological importance of pseurotin in *Penicillium* spp relative to other fungi, we used a dataset of all annotated publicly available genomes originally downloaded from NCBI on April 20th, 2020. This data set comprised 1464 genomes representing 808 species. We performed pairwise reciprocal-best hit analysis of all proteins in the *Aspergillus fumigatus* genome against all 1464 genomes using methods described previously ^75^. Results of this analysis were used to identify *psoA*, *psoB*, *psoC*, *psoD*, *psoE*, *psoF*, *psoG*, and *fapR* orthologs across the fungal kingdom.

We mapped the phylogenetic relationship of fungal genomes based on 290 benchmarking single copy orthologs (BUSCOs) as identified with BUSCO ^76^. We aligned sequences of each BUSCO using MAFFT and trimmed alignments using TrimAl ^77^ with the parameter -automated1. We constructed phylogenies for each gene using IQtree ^78^ after testing for the best fit model. We then created a single consensus tree using ASTRAL ^79^.

We selected a subset of genomes to represent taxonomic breadth based on visual inspection of phylogenetic relationships between species where seven or eight pseurotin genes were found. Additionally, we added a set of genomes that were not present on NCBI but were found to contain the pseurotin gene cluster from our analysis of this cluster in the Penicillium genus (above). Phylogenetic relationships between these genomes were determined using the same methodology as described above for whole-kingdom level analysis. Pseurotin gene clusters were determined from antiSMASH ^80^ by selecting cluster calls that contained the psoA ortholog (as determined above). Resulting clusters were aligned and visualized using clinker ^81^. When BGC border calls made by antiSMASH extended beyond the pseurotin BGC, we trimmed these calls to facilitate visualization of this gene cluster. Validation of gene calls in genome annotations was beyond the scope of this study. To explore the evolutionary history of the pseurotin gene cluster, we aligned PsoA, PsoB, and PsoC sequences of the selected species using MAFFT ^82^ using the same parameters as above. Resulting alignments were trimmed with TrimAl ^77^ then concatenated to form a single sequence for each species. A phylogeny was generated from this alignment using IQtree ^78^. Phylogenies were visualized in R ^83^ using ggtree ^84^.

## DATA AVAILABILITY

Sequence data that support the findings of this study have been deposited in the NCBI SRA database with accession numbers PRJNA861320 for the whole-genome resequencing data and PRJNA861316 for the RNA-seq data. The LC-MS raw data have also been deposited to MassIVE (https://massive.ucsd.edu/ProteoSAFe/static/massive.jsp) and data is available at ftp://massive.ucsd.edu/MSV000090563/ - Dataset ID: MSV000090563, Password: cheese Source data used to create all figures is available in the Supplementary Data provided with the paper.

## AVAILABILITY OF BIOLOGICAL MATERIALS

All unique materials, including the *Penicillium sp.* strain MB WT isolated from cheese, the *Penicillium* sp. strain MB *laeA* deletion mutant, and the bacterial strains used in the interaction and antimicrobial assays, are readily available from the authors upon request.

## ACKNOWLEDGEMENTS

This work was supported by a grant from the National Science Foundation (CAREER # 1942063) to B.W., a Secure Ecosystem Engineering and Design (SEED) project funded by the Genomic Science Program of the U.S. Department of Energy, Office of Science, Office of Biological and Environmental Research (BER) as part of the Secure Biosystems Design Science Focus Area (SFA) to P.E.A.. Ruby Ye provided feedback on this manuscript and experimental design.

## AUTHOR CONTRIBUTIONS

Conceptualization was carried out by J.T., B.E.W., and N.P.K. Experiments were performed by J.T., C.C., M.T.D., and B.E.W. The article was written by J.T., C.C., and B.E.W., and revised with input from all authors. The figures and statistical analyses were made by J.T., C.C., T.A.R., and B.E.W., except for Figure 4 (M.T.D.), Supplementary Figure 10 (R.J.G.), and Supplementary Figure 11 (M.T.D.). The RNA-seq experiments were conducted and analyzed by B.W. The LC-MS curation of data and analysis were conducted by R.J.G, P.E.A., T.A.R. and J.T. The study was supervised by N.P.K. and B.E.W.

## SUPPLEMENTARY FIGURE CAPTIONS

**Supplementary Fig. 1: Deletion and complementation of *laeA* gene in *Penicillium* strain MB.** (a) Schematic representation of the genetic construct for *laeA* deletion in strain MB. The construct is constituted of a gene conferring resistance to hygromycin under TrpC promoter and terminator. (b) Southern blot analyzes of genomic DNA from the WT, the Δ*laeA* and *HygR* strains. The positions of the restriction enzyme cutting sites used for southern blot are shown on the construct schematic. Ten micrograms of total DNA from each strain were digested with the appropriate enzymes and subjected to southern blot analysis using respectively the 5’ flank fragment (blue) and the 3’fragment (grey) as probes. The 1 kb DNA ladder from New England Biolabs was used to determine the size of the expected bands. (c) Restriction map of plasmid pBC-phleo containing the ble gene for phleomycin resistance controlled by the *Aspergillus nidulans* gpdA promoter and the *Saccharomyces cerevisiae* CYC1 terminator. Restriction site used for cloning of the *laeA* gene is shown in red. (d) Southern confirmation of the single insertion of *laeA* using NotI as restriction enzyme. The positive control corresponds to the digested PJT3 plasmid.

**Supplementary Fig. 2: Growth rates of *Penicillium sp.* on YES, PDA, CYA, and CCA.** (a) Growth on YES medium, (b) growth on PDA medium, and (c) growth on CYA medium. In the box plots, the bar represents the standard errors of the means, and each dot represents a biological replication (n=3). One-way analysis of variance (ANOVA) was performed for each bacterial community. Dunnett’s multiple comparison test was used and compared to the WT strain. (****) indicates a *p*-value <0.0001, (***) indicates a *p*-value <0.001, (**) indicates a *p*-value <0.01, (*) indicates a *p*-value <0.05, and no asterisk indicates not significant (ns). For exact *p*-values for each treatment, see **Dataset 1**. (d) Growth on CCA, where points and connecting lines with standard errors of the means bars were used from two independent experiments with five biological replications each. In experiment 2, replication 5 was removed due to no growth of the bacteria in the control. Two-way analysis of variance (ANOVA) was performed for each fungal strain. Dunnett’s multiple comparison test was used and compared to the WT strain. (****) indicates a *p*-value <0.0001, (***) indicates a *p*-value <0.001, (**) indicates a *p*-value <0.01, (*) indicates a *p*-value <0.05, and no asterisk indicates not significant (ns). For exact *p*-values for each treatment, see **Dataset 1.**

**Supplementary Fig. 3: Community analysis in the presence of another common cheese rind fungal community member, the yeast *Debaryomyces hansenii.*** Histograms showing the shifts in a typical rind community (*Staphylococcus, Brevivacterium, Brachybacterium* and *Psychrobacter*) in the presence *Penicillium* sp. strain MB_WT and mutants (Δ*laeA*, Hyg^R^, and *laeAc*) and *D. hansenii* after (a) 3, (c) 10, and (e) 21 days of incubation. Data are mean relative abundance from two independent experiments. Histograms showing the inhibition of community members grown in the presence *Penicillium* sp. strain MB_WT, mutant strains, and *D. hansenii* after (b) 3 (d), 10 and (f) 21 days of incubation. Data are from two independent experiments with five biological replications each. In the box plots, the bar represents the standard errors of the means, and each dot represents a biological replication. One-way analysis of variance (ANOVA) was performed for each bacterial community. Dunnett’s multiple comparison test was used and compared to the WT. (****) indicates a *p*-value <0.0001, (***) indicates a *p*-value <0.001, (**) indicates a *p*-value <0.01, (*) indicates a *p*-value <0.05, and no asterisk indicates not significant (ns). For exact *p*-values for each treatment, see **Dataset 1**.

**Supplementary Fig. 4: Antibacterial activities of filtered crude extracts from cultures on different media.** Crude extracts are collected from cultures of the different *Penicillium* strains on: (a) YES, (b) PDA (c) CYA, and (d) CCA media. The diameter of the zone of inhibition was measured and analyzed. In the box plots, the bar represents the standard errors of the means, and each dot represents a biological replication (n=3). One-way analysis of variance (ANOVA) was performed for each bacterial community. Dunnett’s multiple comparison test was used and compared to the WT strain. (****) indicates a *p*-value <0.0001, (***) indicates a *p*-value <0.001, (**) indicates a *p*-value <0.01, (*) indicates a *p*-value <0.05, and no asterisk indicates not significant (ns). For exact *p*-values for each treatment, see **Dataset 1.**

**Supplementary Fig. 5: Knock out of the *psoA* gene in *Penicillium* strain MB** (a) Map of plasmid pCS01 from which the hygromycin deletion cassette was amplified, used to knock-out, and deletion of psoA gene in Penicillium strain MB. (b) Scheme of the experimental strategy adopted to knockout the *psoA* gene in the *Penicillium* strain MB using CRISPR-Cas9 system. The CRISPR components were delivered to the strain MB_WT protoplasts as a ribonucleoprotein (RNP) complex consisting of the Cas9 nuclease and single guide RNA (sgRNA) targeting the PKS/NRPS conserved domain in the *psoA* gene sequence. The RNP complex will bind to the target sequence at the specific site and cut DNA double strands. The DNA double-strand breaks will be repaired by homology directed repair using the donor DNA template containing 1 kb of homology arms flanking the *hph* gene that confers resistance to hygromycin. (c) Re-sequencing the genomes of the Δ*psoA* strains revealed the deleted regions by the CRISPR-Cas9 mediated knockout. Read mappings for the two different mutants are shown and the deleted regions are represented by the large gap in both mappings. DNA resequencing reads are black lines.

**Supplementary Fig. 6: LC-MS data showing the absence of pseurotin production in the crude extracts of both MB_Δ*psoA* strains compared to the MB_WT strain.** (a) Chromatograms showing the absence of pseurotin A and putative pseurotin D peaks in the Δ*psoA* mutants (b) Fragmentation mass spectrum in positive ionization mode of pseurotin A in MB_WT strain compared to pseurotin A standard (c) Fragmentation mass spectrum in positive ionization mode putative pseurotin D in MB_WT strain.

**Supplementary Figure 7: The frequency of genes associated with the pseurotin gene cluster (left) across fungal phyla (right).** A total of 1463 fungal genomes representing 808 species were obtained from NCBI. Phylogenetic relationships between genomes was determined from the consensus of 290 maximum-likelihood trees constructed for single-copy orthologs (BUSCOs). Terminal branch lengths are not calculated. The number of pseurotin genes present in each genome was determined using reciprocal best-hit BLAST analysis using the *Aspergillus fumigatus* pseurotin genes *psoA, psoB, psoC, psoD, psoE, psoF, psoG,* and *fapR.* The orange arrow indicates a monophyletic clade within Ascomycota that was visually identified as having a higher prevalence of pseurotin genes.

## REFERENCES

1. Fleming, A. On the Antibacterial Action of Cultures of a Penicillium, with Special Reference to their Use in the Isolation of B. influenzæ. Br. J. Exp. Pathol. 10, 226 (1929).

2. Peplow, A. W., Meek, I. B., Wiles, M. C., Phillips, T. D. & Beremand, M. N. Tri16 is required for esterification of position C-8 during trichothecene mycotoxin production by Fusarium sporotrichioides. Appl. Environ. Microbiol. 69, 5935–5940 (2003).

3. Bhatnagar, D., Cary, J. W., Ehrlich, K., Yu, J. & Cleveland, T. E. Understanding the genetics of regulation of aflatoxin production and Aspergillus flavus development. Mycopathologia 162, 155–166 (2006).

4. Georgianna, D. R. & Payne, G. A. Genetic regulation of aflatoxin biosynthesis: from gene to genome. Fungal Genet. Biol. 46, 113–125 (2009).

5. Gil-Serna, J., Vázquez, C. & Patiño, B. Genetic regulation of aflatoxin, ochratoxin A, trichothecene, and fumonisin biosynthesis: A review. Int. Microbiol. 23, 89–96 (2020).

6. Keller, N. P., Turner, G. & Bennett, J. W. Fungal secondary metabolism — from biochemistry to genomics. Nature Reviews Microbiology vol. 3 937–947 Preprint at https://doi.org/10.1038/nrmicro1286 (2005).

7. Betina, V., Mičeková, D. & Nemec, P. Antimicrobial properties of cytochalasins and their alteration of fungal morphology. Microbiology 71, 343–349 (1972).

8. Calvo, A. M., Wilson, R. A., Bok, J. W. & Keller, N. P. Relationship between secondary metabolism and fungal development. Microbiol. Mol. Biol. Rev. 66, 447–59, table of contents (2002).

9. Calvo, A. M. & Cary, J. W. Association of fungal secondary metabolism and sclerotial biology. Front. Microbiol. 6, 62 (2015).

10. Yim, G., Wang, H. H. & Davies, J. Antibiotics as signalling molecules. Philos. Trans. R. Soc. Lond. B Biol. Sci. 362, 1195–1200 (2007).

11. Maillet, F. et al. Fungal lipochitooligosaccharide symbiotic signals in arbuscular mycorrhiza. Nature 469, 58–63 (2011).

12. Pusztahelyi, T., Holb, I. J. & Pócsi, I. Secondary metabolites in fungus-plant interactions. Front. Plant Sci. 6, 573 (2015).

13. Rush, T. A. et al. Lipo-chitooligosaccharides as regulatory signals of fungal growth and development. Nat. Commun. 11, 3897 (2020).

14. Scharf, D. H., Heinekamp, T. & Brakhage, A. A. Human and plant fungal pathogens: the role of secondary metabolites. PLoS Pathog. 10, e1003859 (2014).

15. Snini, S. P. et al. Patulin is a cultivar-dependent aggressiveness factor favouring the colonization of apples by Penicillium expansum. Mol. Plant Pathol. 17, 920–930 (2016).

16. Losada, L., Ajayi, O., Frisvad, J. C., Yu, J. & Nierman, W. C. Effect of competition on the production and activity of secondary metabolites in Aspergillus species. Med. Mycol. 47 **Suppl 1**, S88–96 (2009).

17. Drott, M. T., Debenport, T., Higgins, S. A., Buckley, D. H. & Milgroom, M. G. Fitness Cost of Aflatoxin Production in Aspergillus flavus When Competing with Soil Microbes Could Maintain Balancing Selection. MBio 10, (2019).

18. Costa, J. H., et al. Antifungal potential of secondary metabolites involved in the interaction between citrus pathogens. Scientific Reports vol. 9 Preprint at https://doi.org/10.1038/s41598-019-55204-9 (2019).

19. Rohlfs, M., Albert, M., Keller, N. P. & Kempken, F. Secondary chemicals protect mould from fungivory. Biol. Lett. 3, 523–525 (2007).

20. Drott, M. T., Lazzaro, B. P., Brown, D. L., Carbone, I. & Milgroom, M. G. Balancing selection for aflatoxin in Aspergillus flavus is maintained through interference competition with, and fungivory by insects. Proc. Biol. Sci. 284, (2017).

21. Jakubczyk, D. & Dussart, F. Selected Fungal Natural Products with Antimicrobial Properties. Molecules 25, (2020).

22. Aly, A. H., Debbab, A. & Proksch, P. Fifty years of drug discovery from fungi. Fungal Divers. 50, 3–19 (2011).

23. Keller, N. P. Fungal secondary metabolism: regulation, function and drug discovery. Nat. Rev. Microbiol. 17, 167–180 (2019).

24. Bayram, O. et al. VelB/VeA/LaeA complex coordinates light signal with fungal development and secondary metabolism. Science 320, 1504–1506 (2008).

25. Bok, J. W. & Keller, N. P. LaeA, a regulator of secondary metabolism in Aspergillus spp. Eukaryot. Cell 3, 527–535 (2004).

26. Perrin, R. M. et al. Transcriptional regulation of chemical diversity in Aspergillus fumigatus by LaeA. PLoS Pathog. 3, e50 (2007).

27. Kim, H.-K. et al. Functional roles of FgLaeA in controlling secondary metabolism, sexual development, and virulence in Fusarium graminearum. PLoS One 8, e68441 (2013).

28. Kumar, D. et al. LaeA regulation of secondary metabolism modulates virulence in Penicillium expansum and is mediated by sucrose. Mol. Plant Pathol. (2016) doi:10.1111/mpp.12469.

29. Knowles, S. L. et al. Mapping the Fungal Battlefield: Using in situ Chemistry and Deletion Mutants to Monitor Interspecific Chemical Interactions Between Fungi. Front. Microbiol. 10, 285 (2019).

30. Irlinger, F., Layec, S., Hélinck, S. & Dugat-Bony, E. Cheese rind microbial communities: diversity, composition and origin. FEMS Microbiol. Lett. 362, 1–11 (2015).

31. Ropars, J., Cruaud, C., Lacoste, S. & Dupont, J. A taxonomic and ecological overview of cheese fungi. Int. J. Food Microbiol. 155, 199–210 (2012).

32. Frisvad, J. C. Taxonomy, chemodiversity, and chemoconsistency of Aspergillus, Penicillium, and Talaromyces species. Front. Microbiol. 5, 773 (2014).

33. Diblasi, L., Arrighi, F., Silva, J., Bardón, A. & Cartagena, E. Penicillium commune metabolic profile as a promising source of antipathogenic natural products. Nat. Prod. Res. 29, 2181–2187 (2015).

34. Cabañes, F. J., Bragulat, M. R. & Castellá, G. Ochratoxin A producing species in the genus Penicillium. Toxins 2, 1111–1120 (2010).

35. Casquete, R. et al. Cyclopiazonic acid gene expression as strategy to minimizing mycotoxin contamination in cheese. Fungal Biol. 125, 160–165 (2021).

36. Puel, O., Galtier, P. & Oswald, I. P. Biosynthesis and toxicological effects of patulin. Toxins 2, 613–631 (2010).

37. Martín, J. F. & Liras, P. Secondary Metabolites in Cheese Fungi. in Fungal Metabolites (eds. Mérillon, J.-M. & Ramawat, K. G.) 293–315 (Springer International Publishing, 2017).

38. Vallone, L., Giardini, A. & Soncini, G. Secondary Metabolites from Penicillium roqueforti, A Starter for the Production of Gorgonzola Cheese. Ital J Food Saf 3, 2118 (2014).

39. Wolfe, B. E., Button, J. E., Santarelli, M. & Dutton, R. J. Cheese rind communities provide tractable systems for in situ and in vitro studies of microbial diversity. Cell 158, 422–433 (2014).

40. Kastman, E. K. et al. Biotic interactions shape the ecological distributions of Staphylococcus species. mBio 7: e01157–16. Kumar, R., Jangir, PK, Das, J., Taneja, B. and Sharma (2016).

41. Zhang, Y., Kastman, E. K., Guasto, J. S. & Wolfe, B. E. Fungal networks shape dynamics of bacterial dispersal and community assembly in cheese rind microbiomes. Nat. Commun. 9, 336 (2018).

42. Bodinaku, I., et al. Rapid Phenotypic and Metabolomic Domestication of Wild Penicillium Molds on Cheese. mBio vol. 10 Preprint at https://doi.org/10.1128/mbio.02445-19 (2019).

43. Cosetta, C. M. & Wolfe, B. E. Deconstructing and Reconstructing Cheese Rind Microbiomes for Experiments in Microbial Ecology and Evolution. Curr. Protoc. Microbiol. 56, e95 (2020).

44. Gillot, G. et al. 1-Octanol, a self-inhibitor of spore germination in Penicillium camemberti. Food Microbiol. 57, 1–7 (2016).

45. Osbourn, A. Secondary metabolic gene clusters: evolutionary toolkits for chemical innovation. Trends Genet. 26, 449–457 (2010).

46. Wiemann, P. et al. Prototype of an intertwined secondary-metabolite supercluster. Proc. Natl. Acad. Sci. U. S. A. 110, 17065–17070 (2013).

47. Pinheiro, E. A. A. et al. Antibacterial activity of alkaloids produced by endophytic fungus Aspergillus sp. EJC08 isolated from medical plant Bauhinia guianensis. Nat. Prod. Res. 27, 1633–1638 (2013).

48. Samanidou, V., Michaelidou, K., Kabir, A. & Furton, K. G. Fabric phase sorptive extraction of selected penicillin antibiotic residues from intact milk followed by high performance liquid chromatography with diode array detection. Food Chem. 224, 131–138 (2017).

49. Fisch, K. M. Biosynthesis of natural products by microbial iterative hybrid PKS–NRPS. RSC Adv. 3, 18228–18247 (2013).

50. Breitenstein, W., Chexal, K. K., Mohr, P. & Tamm, C. Pseurotin B, C, D, and E. Further New Metabolites ofPseudeurotium ovalis STOLK. Helv. Chim. Acta 64, 379–388 (1981).

51. Martín, J. F. Key role of LaeA and velvet complex proteins on expression of β-lactam and PR-toxin genes in Penicillium chrysogenum: cross-talk regulation of secondary metabolite pathways. J. Ind. Microbiol. Biotechnol. 44, 525–535 (2017).

52. Lv, Y. et al. Insight into the global regulation of laeA in Aspergillus flavus based on proteomic profiling. Int. J. Food Microbiol. 284, 11–21 (2018).

53. Wang, G., et al. Requirement of LaeA, VeA, and VelB on Asexual Development, Ochratoxin A Biosynthesis, and Fungal Virulence in Aspergillus ochraceus. Front. Microbiol. 10, 2759 (2019).

54. Perlatti, B., Lan, N., Jiang, Y., An, Z. & Bills, G. Identification of Secondary Metabolites from Aspergillus pachycristatus by Untargeted UPLC-ESI-HRMS/MS and Genome Mining. Molecules 25, (2020).

55. Ishikawa, M. et al. Pseurotin A and its analogues as inhibitors of immunoglobuline E production. Bioorg. Med. Chem. Lett. 19, 1457–1460 (2009).

56. Mehedi, Molla, Khondkar & Sultana. Pseurotin A: an antibacterial secondary metabolite from Aspergillus fumigatus. *AJIE*.

57. Hayashi, A. et al. Fumiquinones A and B, nematicidal quinones produced by Aspergillus fumigatus. Biosci. Biotechnol. Biochem. 71, 1697–1702 (2007).

58. Sbaraini, N. et al. Intra-hemocoel injection of pseurotin A from Metarhizium anisopliae, induces dose-dependent reversible paralysis in the Greater Wax Moth (Galleria mellonella). Fungal Genet. Biol. 159, 103675 (2022).

59. Martínez-Luis, S. et al. Antiparasitic and anticancer constituents of the endophytic fungus Aspergillus sp. strain F1544. Nat. Prod. Commun. 7, 165–168 (2012).

60. Maiya, S., Grundmann, A., Li, X., Li, S.-M. & Turner, G. Identification of a hybrid PKS/NRPS required for pseurotin A biosynthesis in the human pathogen Aspergillus fumigatus. Chembiochem 8, 1736–1743 (2007).

61. Bertinetti, B. V., Peña, N. I. & Cabrera, G. M. An antifungal tetrapeptide from the culture of Penicillium canescens. Chem. Biodivers. 6, 1178–1184 (2009).

62. Tsunematsu, Y. et al. Elucidation of pseurotin biosynthetic pathway points to trans- ActingC-methyltransferase: Generation of chemical diversity. Angew. Chem. Weinheim Bergstr. Ger. 126, 8615–8619 (2014).

63. Martín, J. F. & Coton, M. Chapter 12 - Blue Cheese: Microbiota and Fungal Metabolites. in Fermented Foods in Health and Disease Prevention (eds. Frias, J., Martinez-Villaluenga, C. & Peñas, E.) 275–303 (Academic Press, 2017).

64. Kure, C. F. & Skaar, I. The fungal problem in cheese industry. Current Opinion in Food Science 29, 14–19 (2019).

65. Cleary, J. L., Kolachina, S., Wolfe, B. E. & Sanchez, L. M. Coproporphyrin III Produced by the Bacterium Glutamicibacter arilaitensis Binds Zinc and Is Upregulated by Fungi in Cheese Rinds. mSystems 3, (2018).

66. Pierce, E. C. et al. Bacterial–fungal interactions revealed by genome-wide analysis of bacterial mutant fitness. Nature Microbiology vol. 6 87–102 Preprint at https://doi.org/10.1038/s41564-020-00800-z (2021).

67. Bok, J. W. & Keller, N. P. Fast and easy method for construction of plasmid vectors using modified quick-change mutagenesis. Methods Mol. Biol. 944, 163–174 (2012).

68. Niccum, B. A., Kastman, E. K., Kfoury, N., Robbat, A., Jr & Wolfe, B. E. Strain-Level Diversity Impacts Cheese Rind Microbiome Assembly and Function. mSystems 5, (2020).

69. Hassan, M. M., Chowdhury, A. K. & Islam, T. In Silico Analysis of gRNA Secondary Structure to Predict Its Efficacy for Plant Genome Editing. in CRISPR-Cas Methods*: Volume* 2 (eds. Islam, M. T. & Molla, K. A.) 15–22 (Springer US, 2021).

70. Trapnell, C., Pachter, L. & Salzberg, S. L. TopHat: discovering splice junctions with RNA-Seq. Bioinformatics 25, 1105–1111 (2009).

71. Love, M. I., Huber, W. & Anders, S. Moderated estimation of fold change and dispersion for RNA-seq data with DESeq2. Genome Biol. 15, 550 (2014).

72. Xie, C. et al. KOBAS 2.0: a web server for annotation and identification of enriched pathways and diseases. Nucleic Acids Res. 39, W316–22 (2011).

73. Tannous, J. et al. Fungal attack and host defence pathways unveiled in near-avirulent interactions of Penicillium expansum creA mutants on apples. Mol. Plant Pathol. 19, 2635–2650 (2018).

74. MacLean, B., et al. Skyline: an open source document editor for creating and analyzing targeted proteomics experiments. Bioinformatics 26, 966–968 (2010).

75. Drott, M. T., et al. Diversity of Secondary Metabolism in Aspergillus nidulans Clinical Isolates. mSphere vol. 5 Preprint at https://doi.org/10.1128/msphere.00156-20 (2020).

76. Simão, F. A., Waterhouse, R. M., Ioannidis, P., Kriventseva, E. V. & Zdobnov, E. M. BUSCO: assessing genome assembly and annotation completeness with single-copy orthologs. Bioinformatics vol. 31 3210–3212 Preprint at https://doi.org/10.1093/bioinformatics/btv351 (2015).

77. Capella-Gutiérrez, S., Silla-Martínez, J. M. & Gabaldón, T. trimAl: a tool for automated alignment trimming in large-scale phylogenetic analyses. Bioinformatics 25, 1972–1973 (2009).

78. Nguyen, L.-T., Schmidt, H. A., von Haeseler, A. & Minh, B. Q. IQ-TREE: a fast and effective stochastic algorithm for estimating maximum-likelihood phylogenies. Mol. Biol. Evol. 32, 268–274 (2015).

79. Mirarab, S. & Warnow, T. ASTRAL-II: coalescent-based species tree estimation with many hundreds of taxa and thousands of genes. Bioinformatics 31, i44–52 (2015).

80. Blin, K. et al. antiSMASH 5.0: updates to the secondary metabolite genome mining pipeline. Nucleic Acids Res. 47, W81–W87 (2019).

81. Gilchrist, C. L. M. & Chooi, Y.-H. Clinker & clustermap.js: Automatic generation of gene cluster comparison figures. Bioinformatics (2021) doi:10.1093/bioinformatics/btab007.

82. Katoh, K. & Standley, D. M. MAFFT multiple sequence alignment software version 7: improvements in performance and usability. Mol. Biol. Evol. 30, 772–780 (2013).

83. Bunn, A. & Korpela, M. Crossdating in dplR. https://cran.microsoft.com/snapshot/2014-09-08/web/packages/dplR/vignettes/xdate-dplR.pdf (2014).

84. Yu, G., Smith, D. K., Zhu, H., Guan, Y. & Lam, T. T.-Y. Ggtree : An r package for visualization and annotation of phylogenetic trees with their covariates and other associated data. Methods Ecol. Evol. 8, 28–36 (2017).

